# Establishment gaps in species-poor grasslands: artificial biodiversity hotspots to support the colonization of target species

**DOI:** 10.1101/2020.01.23.916155

**Authors:** Réka Kiss, Balázs Deák, Béla Tóthmérész, Tamás Miglécz, Katalin Tóth, Péter Török, Katalin Lukács, Laura Godó, Zsófia Körmöczi, Szilvia Radócz, András Kelemen, Judit Sonkoly, Anita Kirmer, Sabine Tischew, Eva Švamberková, Orsolya Valkó

**Affiliations:** MTA-ÖK Lendület Seed Ecology Research Group, Vácrátót, Hungary; MTA-DE Biodiversity and Ecosystem Services Research Group, Debrecen, Hungary; Department of Ecology, Faculty of Science and Technology, University of Debrecen, Debrecen, Hungary; MTA-DE Lendület Functional and Restoration Ecology Research Group, Debrecen, Hungary; University of Debrecen, Faculty of Science and Technology, Juhász Nagy Pál Doctoral School, Debrecen, Hungary; Anhalt University of Applied Sciences, Bernburg, Germany; Department of Botany, Faculty of Science, University of South Bohemia, České Budějovice, Czech Republic

**Keywords:** colonization, establishment, grazing, grassland restoration, seed sowing, seed mixture

## Abstract

Sowing of grass seed mixtures is a feasible and cost-effective method for landscape-scale grassland restoration. However, sowing only grasses usually leads to species-poor and dense grass sward, where the establishment of target forbs is hampered both by microsite and propagule limitation. To overcome these limitations and increase the diversity of species-poor sown grasslands we developed a novel method by creating ‘establishment gaps’. We used tillage to open gaps of 1 m^2^, 4 m^2^ and 16 m^2^ size in the dense grass sward of six species-poor sown grasslands in the Great Hungarian Plain. We sowed high-diversity seed mixtures of 35 native species into all gaps. We analyzed vegetation development during the first five years after setting up the trial. We also studied the colonization dynamics of the sown species along four 20-m transects around each gap, resulting in a total of 1440 plots of 1 m^2^ size that were studied. Our results indicated that most of the sown species were able to establish permanently in the establishment gaps. The total cover and the cover of perennial sown species increased independently of gap size. Meanwhile the cover of short-lived sown species decreased during the five years. There was only a moderate level of weed abundance in the gaps, and weed cover decreased over the years. The sown target species started to colonize the species-poor grasslands surrounding the establishment gaps within five years. The highest number of species and individuals dispersed from the 4 m^2^-sized gaps, as they had a more stable development than small gaps and were exposed to lower grazing pressure than large ones.

**Implications for practice:**

- Establishment gaps are widely applicable tools to increase the diversity of species-poor grasslands. Gaps of 4 m^2^ represent a more feasible solution compared to larger openings also for the farmers, because there is only a moderate level of weed encroachment and smaller soil disturbance occurs during their creation.
- We recommend sowing high-diversity seed mixtures containing both short-lived species that can establish in the first year and perennial species, which guarantee a high cover of target species later on.
- Gaps sown with high-diversity seed mixture are highly resistant to unfavorable climatic conditions: increasing grass abundance in dry years does not hamper the recovery of target grassland species in the following years.

## Introduction

Grasslands harbor a high diversity of plant and animal species, including endemic and endangered ones (Dengler et al. 2014). Existence of European semi-natural grassland habitats highly depends on traditional management practices (Babai & Molnár 2014; Pruchniewicz 2017); thus, they are threatened by changes in land use, management and disturbance regimes (Helm et al. 2006; Valkó et al. 2018). Cessation of agricultural use in marginal croplands and in some intensively used grasslands give an opportunity for grassland restoration and conservation (Valkó et al. 2016a). Soil seed banks can be a source of species which contribute to the spontaneous recovery of grasslands (Kiss et al. 2016); however, its restoration potential is limited because there are lot of grassland species, which have no persistent seed bank (Bossuyt & Honnay 2008; Valkó et al. 2011). Therefore, in severely degraded areas we cannot rely on spontaneous grassland recovery solely from the seed bank (Klaus et al. 2018). This propagule limitation can be aggravated by the lack of seed rain of target grassland species due to their low dispersal ability and the lack of source populations in intensively used landscapes (Buisson et al. 2006; Novák & Konvička 2006; Deák et al. 2018). Besides propagule limitation, favorable niches for the establishment of grassland species are also limited in perennial-dominated, dense grassland swards. Both the accumulation of living and dead biomass and the encroachment of competitor species can hinder germination and establishment of grassland species (Köhler et al. 2005; Valkó et al. 2016a). Propagule and microsite limitation together halt the spontaneous recovery of species-rich grasslands and their effect generally increases with the time elapsed since the beginning of the degradation (Valkó et al. 2018).

The success of restoration activities can be enhanced by overcoming the two major limitations, by propagule addition to the target area (e.g. seed sowing, hay and topsoil transfer; Kiehl et al. 2010; Török et al. 2011) and by provision of microsites either by natural (animal perturbations or grazing by wild animals) or human-induced (mowing, tilling, grazing by livestock) disturbances (Bullock et al. 1995; Coiffait-Gombault et al. 2012; Valkó et al. 2016b). Such gaps are competition free habitat patches which are important especially in the early and most vulnerable stages of plant establishment (Silwertown & Smith 1988; Grime 2001; Hölzel 2005). Small sized gaps often face the problem of fast recolonization by vegetative spreading species, in larger gaps establishment from soil seed bank, seed rain or sowing is possible (Bullock et al. 1995; Pywell et al. 2007; Eckstein et al. 2012).

In species poor communities gaps can become species rich islets increasing the diversity of the grassland. Pywell et al. (2007) used this approach to create species rich patches by deturfing in grasslands for supporting the colonization of target species, while Benayas et al. (2008) suggested the use of woodland islets to enhance woodland development in former agricultural lands. The low cost of creating these patches, the small area used for restoration purposes and the concentrated high density of target species in patches, which can colonize the surrounding species-poor areas are attractive not only to restoration ecologists but also to the farmers (Benayas et al 2008).

The idea of using ‘establishment gaps’ for increasing the plant diversity of species-poor grasslands is based on decreasing the microsite limitation by opening gaps in the sward and decreasing propagule limitation by sowing seeds of target species in the openings (Valkó et al. 2016b). This method have been developed and tested in restored dry grasslands in Hungary, and the vegetation development in the first two years after gap creation suggested that the method is feasible for introducing target species inside the gaps (Valkó et al. 2016b).

In this study we followed the vegetation development inside the establishment gaps for five years and also tested whether the introduced target species are able to colonize the surrounding grasslands from the gaps. We tested the effectiveness of creating establishment gaps of different sizes with subsequent sowing of 35 target grassland species (sown species) on the diversification of species-poor grasslands. We monitored the establishment success of the 35 sown species in the establishment gaps five years after sowing, and analyzed the dynamics of their colonization of the surrounding species-poor grasslands. We asked the following questions: (i) Which gap size is most favorable for the establishment and colonization of sown species? (ii) Which sown species are most successful in establishment and colonization? (iii) What is the spatial and temporal dynamics of the sown species over the five study years?

## Materials and methods

### Study sites

The study sites are situated in the Hortobágy National Park, East-Hungary, near the towns Egyek and Tiszafüred, at an elevation of 88-92 m a.s.l. (Valkó et al. 2016b). The climate is continental with a mean annual precipitation of 550 mm and mean annual temperature of 9.5 °C with high inter-annual fluctuations (Lukács et al. 2015). In historical times, alkaline marshes (*Bolboschoenetalia maritimi*), alkaline meadows (*Beckmannion eruciformis*), alkaline dry grasslands (*Artemisio-Festucetalia pseudovinae*) and loess grasslands (*Festucion rupicolae*) were typical in the study region (Deák et al. 2014a,b). In the last century, most of the rather fertile loess grasslands and many alkaline grasslands were converted into arable land for crop production (Török et al. 2010). After the abandonment of croplands, the area was sown by low-diversity grass seed mixtures in 2005 (Török et al. 2010). Although the area of the grasslands has been increased by this project, most of the sown grasslands remained species-poor, as the establishment of target grassland species was hampered by propagule and microsite limitation.

### Establishment gaps

To increase the diversity of the species-poor sown grasslands, we created establishment gaps at six sown grassland sites in October 2013 as described by Valkó et al. (2016b). After soil preparation by digging, rotary hoeing and raking, a high-diversity seed mixture, composed of 35 target grassland species (for the species list, see Table S1), was sown in three establishment gaps per site in a density of 10 g/m^2^ (Figure S1). Target grassland species were selected from the regional species pool of loess and alkaline dry grasslands in consultation with the experts of the Hortobágy National Park Directorate. We aimed to cover a broad range of species of the two habitat types and to include both common and regionally rare species in the seed mixture. We applied three gap sizes in each site: (i) 1 m^2^ small gaps, (ii) 4 m^2^ medium-sized gaps and (iii) 16 m^2^ large gaps. The establishment gaps were placed at least 50 m apart from each other to avoid accidental propagule exchange among them. Already from the first year on, establishment gaps were extensively grazed by cattle with a stocking rate of 0.5 livestock unit per hectare between April and October.

### Sampling design

We recorded total vegetation cover and percentage cover of all species found in establishment gaps in late June from 2014 to 2018. Grasses sown in the landscape-scale restoration project in 2005 (i.e., *Bromus inermis, Festuca pseudovina, F. rupicola, Poa angustifolia*) were considered as matrix grasses (Valkó et al. 2016b). Spontaneously established adventive competitors (e.g., *Stenactis annua*), ruderal competitors (e.g., *Cirsium arvense*) and agricultural weeds (e.g., *Convolvulus arvensis*) were cumulated to the category “weeds”, based on the social behavior type system of Borhidi (1995). From 2016, we monitored the spatial and temporal colonization dynamics of the sown species in the surrounding species-poor grasslands. Four 20 m transects (containing twenty 1 m × 1 m plots, so-called ‘colonization plots’) running towards the four main directions from each establishment gap were designated (Figure S1). Transects comprised in total 1440 plots of 1 m^2^ size. We recorded the number of sown species (flowering species) and flowering shoots of each sown species in the colonization plots in the three consecutive years (2016-2018). Nomenclature follows Király et al. (2009).

### Statistical analyses

We used generalized linear mixed models (GLMM, Zuur et al. 2009) to analyze the vegetation dynamics inside the establishment gaps. We used gap size (1 m^2^, 4 m^2^, 16 m^2^) and year (2014-2018) as fixed factors, and site identity as a random factor. Dependent variables were: total cover and the cover of sown species, perennial sown species, short-lived sown species, matrix grasses, weeds, perennial weeds and short-lived weeds.

In the GLMM analyses of the colonization dynamics of the sown species, we tested the effect of establishment gap size (1 m^2^, 4 m^2^, 16 m^2^), distance from establishment gap (1-20 m), and year (2016-2018) as fixed factors on the dependent variables (species number, number of individuals, number of flowering species, number of flowering shoots, and the number of individuals of the nine most abundant sown species separately), with site identity as random factor. The GLMMs were run in SPSS 22.0.

## Results

### Vegetation dynamics inside the establishment gaps

During the five study years, we recorded 172 species in the establishment gaps (please see the total species list in Table S1). All sown species were able to establish in at least one of the establishment gaps (Table S1). The total vegetation cover increased in the establishment gaps over the study period (*p* < 0.001, F = 8.649), parallel with the increasing cover of matrix grasses (*p* < 0.001, F = 9.870) and sown species (*p* < 0.001, F = 7.971) (Figure 1). There was a steady increase in the cover of sown species over the study period, with the exception of the year 2017, when their cover decreased considerably but increased again in 2018 (Figure 1). The cover of perennial sown species increased with time (*p* < 0.001, F = 14.587), while the cover of short-lived sown species had a decreasing tendency in all gap sizes (*p* = 0.001, F = 5.499) (Table S2). Matrix grass species reached the highest cover in the last two years (Figure 1). We recorded a total of 57 weed species. The cover of both short-lived and perennial weeds decreased steadily from 2014 until 2018 (*p* < 0.001, F = 9.189) (Figure 1, Table S2).

**Figure 1.**
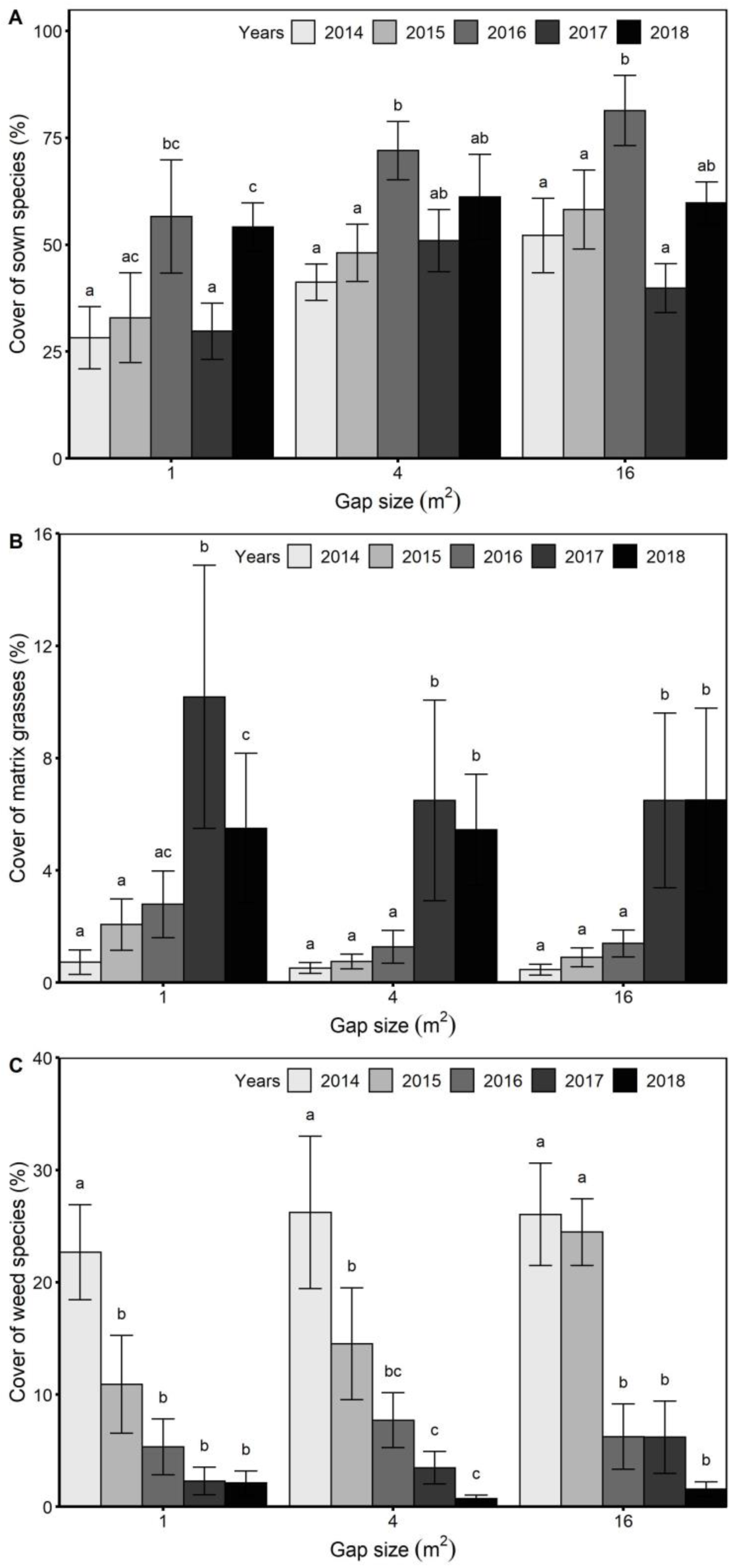
Cover scores (mean ± SE) of sown-, matrix- and weed species in the establishment gaps from 2014 to 2018. Letters indicate significant differences in the cover scores between the years within different gap sizes (GLMM and LSD test, p < 0.05).

### Colonization of sown species from the gaps to the surroundings

#### Temporal dynamics

In total we found 30 out of 35 sown species along the transects, from which 27 species occurred in 2016, 26 in 2017 and 28 in 2018. We recorded in total 38,446 individuals of sown species in the colonization plots in the surroundings of the gaps. The colonization plots hosted on average 10.7 individuals of sown species per m^2^ in 2016, 6.7 in 2017 and 9.2 in 2018, differences between years were significant in every case (*p* < 0.001, F = 340.97) (Figure 2, Table S3). *Achillea collina* and *Centaurea solstitialis* were represented with high individual number in each year; besides them in 2016 *Trifolium striatum* and *Dianthus pontederae* were the most abundant species. In 2017 and 2018 *Trifolium campestre* was present in large numbers (Table S4).

**Figure 2.**
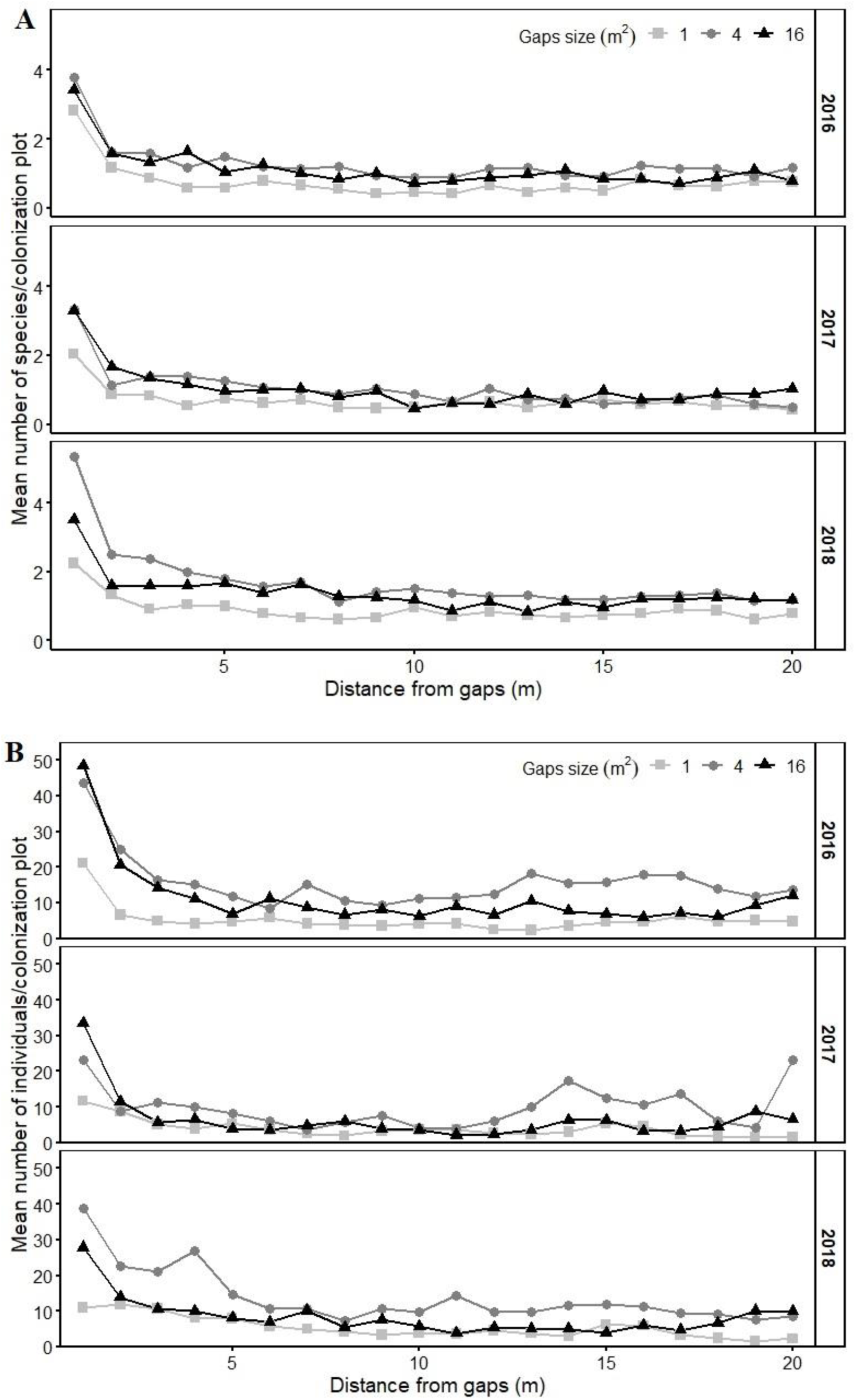
Number of sown species (A) and individuals (B) in the 1 m^2^-sized colonization plots adjacent to different sized (1 m^2^, 4 m^2^, 16 m^2^) establishment gaps in the three study years (2016-2018).

Out of the 30 species, which were found in the colonization plots, 28 species flowered, 23 in the first year, 22 in the second year and 25 in the third year. Over the study period we counted in total 37,849 flowering shoots; the highest number was observed in 2018 (total: 17,488; 12.14/colonization plot), it was lower in 2016 (total: 14,044; mean: 9.75/colonization plot) and the lowest in 2017 (total: 6,317; mean: 4.38/colonization plot). Species numbers and individual numbers of sown species (Figure S2), number of flowering sown species and the number of flowering shoots (Figure S3) were the lowest in 2017. However, all variables, except total number of individuals, reached their highest scores in 2018 (Figure S2, S3).

#### Spatial dynamics

All sown species were present in at least one colonization plot per transect. The species (*p* < 0.001, F = 366.593) and individual numbers of sown species (*p* < 0.001, F = 3052.579) was the highest in the first plots adjacent to the establishment gaps in every study year (Figure 2) and decreased with increasing distance from establishment gaps (Table S3). In the first plots adjacent to the establishment gaps, we found in total 26 sown species in the three study years. The individual number of sown species was more than two-fold higher in the first (in total 6,225 individuals, more than 16% of the total individuals) plot adjacent to the establishment gaps than in any other plots and decreased with increasing distance (Figure 2). The average number of flowering species (*p* < 0.001, F = 176.935) and the number of flowering shoots (*p* < 0.001, F = 3649.264) showed similar dynamics (Table S3, Figure S3).

#### Effect of establishment gaps size on the colonization success of sown species

Gap size had a significant effect on all studied variables (Table S3). The highest number of sown species colonized the adjacent species-poor grassland from the 4 m^2^ medium-sized establishment gaps (*p* < 0.001, F = 44.342) (in total 28 species), while a total of 24 species were present near the 1 m^2^ small establishment gaps and 19 near the 16 m^2^ large gaps. The number of species per plot was highest close to the medium-sized gaps exceeding the numbers of species close to the smallest gaps (Figure 2). We found in total 18,923 individuals in the colonization plots nearest to the medium-sized gaps, 12,479 near the large gaps and 7,044 near the small gaps (*p* < 0.001, F = 360.887). Similar trends were detected for the number of flowering species (*p* < 0.001, F = 29.214) and the number of flowering shoots (*p* < 0.001, F = 324.912) (Figure S4).

## Discussion

### Vegetation development inside the establishment gaps

We demonstrated that soil disturbance with subsequent seed sowing increase plant diversity in species poor grasslands, and that establishment gaps were suitable for increasing diversity over at least a five-year time period (see also Klaus et al. 2017; Valkó et al. 2016b), and presumably even on a longer run. The cover of target grassland species, including perennial sown species and matrix grasses re-growing after disturbance, increased over the studied time period in the establishment gaps. It is especially important that the cover of weed species decreased significantly over the years, as a result of the successful weed suppression effect of the sown species (Tracy & Sanderson 2004; Valkó et al. 2016b; Švamberková et al. 2019). The trends were similar in all gap sizes; however, we found that sown perennial species reached higher cover scores in large gaps than in small ones.

Our results also highlighted the potential of species-rich establishment gaps to overcome stochastic events, such as extreme drought. The year 2017 was a particularly dry year and had significant negative effects on the establishment and colonization of the sown species. The extremely dry winter of 2016/2017 and spring-summer period of 2017 could be the reasons for the low cover scores of the sown species in 2017. There was only 290 mm of precipitation between December 2016 and July 2017 in the study region; while the average rainfall in that period was 370 mm in the studied five years (data from the Hungarian Meteorological Service). Drought likely resulted in the decrease of sown species, consequently opening new gaps for the establishment and encroachment of matrix grasses in the gaps in 2017. Matrix grasses have dense and deep root systems (Kutschera et al. 1982), which gives them a competitive advantage over the sown target species under dry weather conditions. However, the increased cover of matrix grasses did not have a strong effect on the sown species in the long run, because sown species were able to recover already in 2018. The higher species diversity of gaps probably goes with a higher community stability compared with the species poor sown grasslands (portfolio effect; Doak et al. 1998, Tilman et al. 1998). Greater species richness promotes stability of communities (Tilman & Downing 1994), because there are high number of species that respond differently to the environmental fluctuations, so the decline of one of them could be compensated by the strengthening of another one (Ives et al. 2000, Lepš 2004, Polley et al. 2013).

### Temporal and spatial colonization dynamics

The high species diversity of the establishment gaps proved to be the source of the species colonization to the adjacent grasslands. Previous studies reported moderate establishment rate of sown species in the adjacent grasslands (Ruprecht 2006; Albert et al. 2014); therefore, we assume that also in our case it would take several years for grassland species to permanently establish outside the gaps (Baasch et al. 2016, Švamberková et al. 2019). We found species with good dispersal ability and good competitive ability to establish in the adjacent species-poor and dense grassland sward. Out of the nine most successfully established sown species that were well represented in the adjacent grasslands six were most abundant in the proximity of the large- and medium-sized gaps (*Achillea collina, Centaurea solstitialis, Galium verum, Podospermum canum, Potentilla argentea* and *Trifolium campestre*), while the colonization success of the remaining three species was not affected by the gap size. We observed a decline of the *A. collina* individual numbers during the study years, which can be linked to the increased abundance of *C. solstitialis*. Being a highly competitive species with good defensive mechanisms against grazing and high seed production rate, *C. solstitialis* was able to successfully establish and increase its abundance in the gaps and in the surrounding grassland (Callaway et al. 2006; Wallace et al. 2008). *C. solstitialis* is a legally protected plant species in Hungary, however it is being considered as a pest species in many other countries. Its current abundance in the study site does not decrease considerably the forage quality, but its presence might be problematic in the future. The dynamics of *Podospermum canum* is strongly linked to the dynamics of *C. solstitialis.* The increase of *P. canum* might be related to the enhanced presence of the above-mentioned species, which provides physical defense against grazers but it does not hamper the development of *P. canum* due to their different growth forms.

### Effect of establishment gap size on the colonization success of sown species

Small gaps were less effective in supporting establishment and colonization of sown species than the large and medium-sized ones. However, contrary to our expectations, not the largest establishment gaps were the most effective but the medium-sized ones. The 4 m^2^ medium-sized gaps supported the highest species number per colonization plot, the highest number of individuals, the highest number of flowering species, and the highest total number of flowering shoots. Valkó et al. (2016b) found that vegetation development and establishment of sown species is rather stochastic in the small 1 m^2^ gaps. Recolonization by grasses is the most likely in the small gaps with low surface:perimeter ratio, which hinders the establishment and colonization of sown species. Thus, larger gaps are more stable and less susceptible to grass recolonization and provide more favorable conditions for sown species to persist. However, the gaps represent an attractive foraging place for livestock as for their diet they require not only grasses but also forbs having higher carbon:nitrogen ratio (Rutter et al. 2000, Soder et al. 2007). Due to the high visibility of a larger patch of sown species in the otherwise monotone grassland, the larger gaps might have a high probability to be discovered and grazed by animals (Díaz et al. 2001). Grazing animals search for the best tradeoff between intake quantity and quality; therefore, they will repeatedly forage at the re-growing vegetation of large gaps, where grazing occurred previously (Soder et al. 2007). Grazed individuals of target species have a lower seed production rate because they use more energy for survival and re-growth than for seed production (O’Connor & Pickett 1992). As a large gap of 16 m^2^ is more visible than a medium sized gap of 4 m^2^, we suggest that a medium gap size is the best option for grassland diversification. This is confirmed by the highest total number of flowering shoots, which was the largest in the 4 m^2^ gaps. The higher grazing pressure in the largest gaps might thus have resulted in a higher number of flowering shoots per individual, caused mostly by the species like *Centaurea solstitialis*, which is known to not only effectively avoid but also effectively compensate the negative effects of grazing, by compensating lost biomass by increasing bud numbers and seed production rate (Callaway et al. 2006, Wallace et al. 2008).

## Conclusions

Observing the dynamics in establishment gaps for five years highlighted the efficiency of medium-sized establishment gaps (4 m^2^) sown with target grassland species, to increase the diversity of species-poor grasslands. We found that the level of weed encroachment remained low on the created openings already in the first year after sowing and further decreased in parallel with the increasing cover of sown species and matrix grasses. Establishment gaps proved to be an effective source for the colonization of the adjacent species-poor grasslands by target species. Establishment gaps combined with cattle grazing efficiently decrease microsite and propagule limitation, the two main obstacles hampering the establishment of target species in sown grasslands. Our findings suggest that medium-sized gaps can ensure the persistence of sown target species because this gap size is less attractive for grazing animals compared to large gaps; therefore the target species have higher chance for setting seeds. Medium-sized gaps represent the most cost-effective and feasible solution compared to larger openings also for the farmers, who preferred smaller disturbances and a moderate level of weed encroachment.

## Acknowledgments

The authors were supported by NKFI FK 124404 (OV), NKFI KH 126476 (OV), NKFI K 116639 (BT), NKFI KH 126477 (BT), NKFI KH 130338 (BD), NKFI PD 124548 (TM), NKFI PD 128302 (KT), NKFI K119225 (PT), NKFI KH 129483 (PT) and MTA’s Post Doctoral Research Program (AK). OV, BD and AK were supported by the Bolyai János Research Scholarship of the Hungarian Academy of Sciences. OV, BD, AK and KL were supported by the New National Excellence Program of the Hungarian Ministry of Human Capacities. Establishment gaps were created with the support of a project of the German Federal Environmental Foundation (DBU) “Large-scale grassland restoration: the use of establishment gaps and high diversity seeding by the knowledge transfer of regional seed propagation to Hungary (ProSeed)”.

