## Supplemental Table S1-S4, Figure S1-S4 for "Establishment gaps in species-poor grasslands: artificial biodiversity hotspots to support the colonization of target species"

**Table S1. Percentage** cover of species (mean ± SE) in the establishment gaps in the five years. Notations: S – sown target species, W – weeds.

| Year | | 2014 | 2014 | 2014 | 2015 | 2015 | 2015 | 2016 | 2016 | 2016 | 2017 | 2017 | 2017 | 2018 | 2018 | 2018 |
|---|---|---|---|---|---|---|---|---|---|---|---|---|---|---|---|---|
| Gap size (m$^2$) | | 1 | 4 | 16 | 1 | 4 | 16 | 1 | 4 | 16 | 1 | 4 | 16 | 1 | 4 | 16 |
| | S/W | Mean ± SD | Mean ± SD | Mean ± SD | Mean ± SD | Mean ± SD | Mean ± SD | Mean ± SD | Mean ± SD | Mean ± SD | Mean ± SD | Mean ± SD | Mean ± SD | Mean ± SD | Mean ± SD | Mean ± SD |
| *Achillea collina* | S | 2.83 ± 5.52 | 8.25 ± 8.24 | 8.50 ± 6.44 | 5.43 ± 8.06 | 6.08 ± 3.98 | 17.05 ± 11.75 | 4.75 ± 4.36 | 14.17 ± 8.80 | 34.50 ± 13.98 | 9.58 ± 11.41 | 10.08 ± 8.21 | 15.33 ± 13.59 | 5.80 ± 5.94 | 8.78 ± 8.60 | 13.38 ± 11.69 |
| *Aegilops cylindrica* | S | 1.83 ± 0.98 | 2.25 ± 3.43 | 2.45 ± 2.85 | 0.15 ± 0.16 | 0.42 ± 0.78 | 1.00 ± 1.01 | 0.12 ± 0.29 | 0.00 ± 0.00 | 0.33 ± 0.82 | 0.00 ± 0.00 | 0.00 ± 0.00 | 0.00 ± 0.00 | 0.00 ± 0.00 | 0.00 ± 0.00 | 0.00 ± 0.00 |
| *Agrimonia eupatoria* | S | 0.08 ± 0.20 | 0.15 ± 0.21 | 0.23 ± 0.10 | 0.22 ± 0.40 | 0.48 ± 0.26 | 0.57 ± 0.41 | 0.42 ± 0.40 | 1.42 ± 1.25 | 0.98 ± 1.07 | 0.67 ± 0.61 | 0.90 ± 0.86 | 0.42 ± 0.47 | 0.50 ± 0.63 | 1.03 ± 0.53 | 0.95 ± 1.07 |
| *Agropyron cristatum* | S | 0.17 ± 0.41 | 0.42 ± 0.80 | 0.27 ± 0.29 | 0.68 ± 0.96 | 0.53 ± 0.35 | 0.80 ± 1.12 | 0.33 ± 0.82 | 0.05 ± 0.12 | 0.05 ± 0.12 | 0.00 ± 0.00 | 0.27 ± 0.61 | 0.07 ± 0.10 | 0.00 ± 0.00 | 0.00 ± 0.00 | 0.67 ± 0.88 |
| *Allium scorodoprasum* | S | 0.00 ± 0.00 | 0.00 ± 0.00 | 0.00 ± 0.00 | 0.07 ± 0.05 | 0.05 ± 0.12 | 0.05 ± 0.05 | 0.00 ± 0.00 | 0.00 ± 0.00 | 0.18 ± 0.40 | 0.03 ± 0.08 | 0.02 ± 0.04 | 0.05 ± 0.08 | 0.05 ± 0.12 | 0.27 ± 0.61 | 0.02 ± 0.04 |
| *Aster tripolium* | S | 0.00 ± 0.00 | 0.00 ± 0.00 | 0.00 ± 0.00 | 0.00 ± 0.00 | 0.00 ± 0.00 | 0.02 ± 0.04 | 0.00 ± 0.00 | 0.00 ± 0.00 | 0.00 ± 0.00 | 0.00 ± 0.00 | 0.00 ± 0.00 | 0.00 ± 0.00 | 0.00 ± 0.00 | 0.00 ± 0.00 | 0.00 ± 0.00 |
| *Atriplex litoralis* | S | 0.17 ± 0.29 | 0.82 ± 1.10 | 0.65 ± 0.67 | 0.05 ± 0.12 | 0.02 ± 0.04 | 0.00 ± 0.00 | 0.00 ± 0.00 | 0.00 ± 0.00 | 0.00 ± 0.00 | 0.00 ± 0.00 | 0.00 ± 0.00 | 0.00 ± 0.00 | 0.00 ± 0.00 | 0.00 ± 0.00 | 0.00 ± 0.00 |
| *Atriplex tatarica* | S | 0.10 ± 0.20 | 0.17 ± 0.29 | 0.40 ± 0.45 | 0.00 ± 0.00 | 0.00 ± 0.00 | 0.00 ± 0.00 | 0.00 ± 0.00 | 0.00 ± 0.00 | 0.00 ± 0.00 | 0.00 ± 0.00 | 0.00 ± 0.00 | 0.00 ± 0.00 | 0.00 ± 0.00 | 0.00 ± 0.00 | 0.00 ± 0.00 |
| *Bunias orientalis* | S | 0.33 ± 0.43 | 0.12 ± 0.29 | 0.50 ± 0.77 | 0.00 ± 0.00 | 0.00 ± 0.00 | 1.33 ± 2.42 | 0.00 ± 0.00 | 0.67 ± 1.63 | 0.80 ± 1.59 | 0.00 ± 0.00 | 0.03 ± 0.08 | 0.47 ± 1.00 | 0.00 ± 0.00 | 0.43 ± 0.56 | 1.00 ± 2.00 |
| *Bupleurum tenuissimum* | S | 0.00 ± 0.00 | 0.00 ± 0.00 | 0.02 ± 0.04 | 0.00 ± 0.00 | 0.00 ± 0.00 | 0.00 ± 0.00 | 0.00 ± 0.00 | 0.00 ± 0.00 | 0.00 ± 0.00 | 0.02 ± 0.04 | 0.02 ± 0.04 | 0.02 ± 0.04 | 0.00 ± 0.00 | 0.05 ± 0.12 | 0.00 ± 0.00 |
| *Carthamus lanatus* | S | 0.08 ± 0.20 | 0.12 ± 0.29 | 0.17 ± 0.41 | 0.00 ± 0.00 | 0.10 ± 0.15 | 0.65 ± 0.23 | 0.17 ± 0.41 | 0.00 ± 0.00 | 0.22 ± 0.39 | 0.00 ± 0.00 | 0.00 ± 0.00 | 0.03 ± 0.05 | 0.02 ± 0.04 | 0.22 ± 0.40 | 0.05 ± 0.05 |
| *Centaurea jacea angustifolia* | S | 0.97 ± 1.60 | 1.45 ± 2.09 | 2.68 ± 3.80 | 1.60 ± 1.54 | 1.23 ± 0.61 | 2.08 ± 1.36 | 6.58 ± 5.85 | 8.67 ± 7.42 | 5.58 ± 7.12 | 3.92 ± 2.73 | 3.08 ± 2.58 | 5.58 ± 5.46 | 4.08 ± 3.56 | 2.33 ± 1.17 | 3.42 ± 3.53 |
| *Centaurea scabiosa sadleriana* | S | 0.17 ± 0.41 | 1.42 ± 1.56 | 1.00 ± 0.85 | 0.67 ± 1.21 | 1.28 ± 1.52 | 3.83 ± 3.06 | 2.83 ± 3.08 | 6.42 ± 5.89 | 7.50 ± 7.12 | 1.42 ± 1.56 | 4.08 ± 2.25 | 3.33 ± 2.50 | 2.30 ± 2.22 | 5.00 ± 5.22 | 4.17 ± 2.46 |
| *Centaurea solstitialis* | S | 16.25 ± 17.16 | 18.83 ± 10.76 | 25.83 ± 27.92 | 10.83 ± 12.45 | 21.05 ± 23.50 | 12.78 ± 20.85 | 10.5 ± 20.50 | 8.17 ± 9.00 | 5.83 ± 3.25 | 2.00 ± 3.05 | 8.53 ± 14.78 | 3.53 ± 5.76 | 0.30 ± 0.60 | 0.70 ± 0.87 | 1.63 ± 2.42 |
| *Dianthus pontederae* | S | 0.02 ± 0.04 | 0.08 ± 0.20 | 0.13 ± 0.28 | 0.57 ± 0.80 | 1.97 ± 1.74 | 3.30 ± 4.61 | 0.87 ± 1.80 | 0.03 ± 0.05 | 1.37 ± 3.25 | 0.18 ± 0.27 | 0.48 ± 0.62 | 0.10 ± 0.13 | 1.38 ± 2.13 | 1.17 ± 1.03 | 1.80 ± 3.55 |
| *Falcaria vulgaris* | S | 0.02 ± 0.04 | 0.02 ± 0.04 | 0.05 ± 0.12 | 0.25 ± 0.61 | 0.05 ± 0.12 | 0.17 ± 0.15 | 0.25 ± 0.42 | 0.05 ± 0.12 | 0.27 ± 0.39 | 0.05 ± 0.12 | 0.05 ± 0.12 | 0.07 ± 0.08 | 0.07 ± 0.12 | 0.20 ± 0.20 | 0.07 ± 0.12 |
| *Filipendula vulgaris* | S | 0.00 ± 0.00 | 0.03 ± 0.05 | 0.00 ± 0.00 | 0.00 ± 0.00 | 0.10 ± 0.20 | 0.00 ± 0.00 | 0.40 ± 0.79 | 0.08 ± 0.13 | 0.08 ± 0.12 | 0.32 ± 0.59 | 0.20 ± 0.40 | 0.02 ± 0.04 | 0.22 ± 0.40 | 0.60 ± 0.78 | 0.35 ± 0.37 |
| *Galium verum* | S | 0.63 ± 1.18 | 1.33 ± 1.88 | 1.97 ± 2.63 | 2.93 ± 4.60 | 3.67 ± 2.73 | 4.33 ± 1.97 | 5.00 ± 5.54 | 12.08 ± 13.13 | 5.42 ± 4.74 | 3.83 ± 3.66 | 9.67 ± 7.37 | 4.75 ± 5.21 | 14.5 ± 13.49 | 15 ± 14.41 | 13.83 ± 10.83 |
| *Hypericum perforatum* | S | 0.00 ± 0.00 | 0.07 ± 0.12 | 0.13 ± 0.28 | 0.67 ± 1.63 | 0.42 ± 0.80 | 0.22 ± 0.40 | 0.75 ± 1.60 | 1.33 ± 2.42 | 0.43 ± 0.65 | 0.67 ± 1.63 | 0.02 ± 0.04 | 0.08 ± 0.10 | 0.92 ± 1.63 | 0.17 ± 0.29 | 0.27 ± 0.37 |
| *Lathyrus hirsutus* | S | 0.00 ± 0.00 | 0.00 ± 0.00 | 0.05 ± 0.12 | 0.30 ± 0.6 | 0.08 ± 0.20 | 0.08 ± 0.20 | 0.00 ± 0.00 | 0.00 ± 0.00 | 0.05 ± 0.12 | 0.00 ± 0.00 | 0.00 ± 0.00 | 0.00 ± 0.00 | 0.08 ± 0.20 | 0.67 ± 0.75 | 0.08 ± 0.20 |
| *Lathyrus tuberosus* | S | 0.12 ± 0.29 | 0.00 ± 0.00 | 0.08 ± 0.20 | 1.55 ± 2.73 | 0.20 ± 0.32 | 0.25 ± 0.42 | 0.88 ± 1.58 | 0.17 ± 0.29 | 0.03 ± 0.08 | 0.22 ± 0.40 | 0.25 ± 0.61 | 0.02 ± 0.04 | 0.75 ± 1.17 | 0.42 ± 0.58 | 0.23 ± 0.39 |
| *Lotus corniculatus* | S | 2.13 ± 2.79 | 1.83 ± 2.14 | 1.97 ± 1.80 | 2.22 ± 4.81 | 2.63 ± 3.79 | 2.43 ± 3.12 | 2.02 ± 3.00 | 5.08 ± 5.00 | 2.68 ± 3.26 | 0.50 ± 1.22 | 4.42 ± 5.80 | 0.2 ± 0.19 | 0.75 ± 0.99 | 5.08 ± 7.35 | 0.72 ± 0.71 |
| *Plantago media* | S | 0.08 ± 0.20 | 0.28 ± 0.45 | 0.33 ± 0.34 | 0.50 ± 0.84 | 1.17 ± 1.47 | 1.68 ± 1.87 | 0.67 ± 1.21 | 1.22 ± 0.74 | 1.93 ± 1.47 | 0.58 ± 1.19 | 0.33 ± 0.58 | 0.38 ± 0.30 | 1.35 ± 2.34 | 0.95 ± 0.72 | 0.97 ± 0.89 |
| *Podospermum canum* | S | 0.05 ± 0.12 | 0.08 ± 0.20 | 0.22 ± 0.28 | 0.13 ± 0.22 | 0.50 ± 0.36 | 0.48 ± 0.55 | 0.05 ± 0.12 | 1.00 ± 2.45 | 0.42 ± 0.60 | 0.02 ± 0.04 | 0.02 ± 0.04 | 0.07 ± 0.10 | 0.13 ± 0.22 | 0.07 ± 0.12 | 0.07 ± 0.12 |
| *Potentilla argentea* | S | 0.13 ± 0.22 | 0.23 ± 0.39 | 0.20 ± 0.11 | 0.42 ± 1.02 | 0.33 ± 0.43 | 0.65 ± 0.75 | 1.33 ± 2.36 | 1.13 ± 1.31 | 2.50 ± 2.95 | 0.20 ± 0.4 | 0.42 ± 0.56 | 0.17 ± 0.08 | 0.63 ± 1.19 | 1.95 ± 1.43 | 0.7 ± 0.71 |
| *Rapistrum perenne* | S | 0.00 ± 0.00 | 0.00 ± 0.00 | 0.00 ± 0.00 | 0.00 ± 0.00 | 0.05 ± 0.12 | 0.00 ± 0.00 | 0.00 ± 0.00 | 0.00 ± 0.00 | 0.17 ± 0.41 | 0.00 ± 0.00 | 0.00 ± 0.00 | 0.00 ± 0.00 | 0.00 ± 0.00 | 0.00 ± 0.00 | 0.42 ± 1.02 |
| *Salvia verticillata* | S | 0.00 ± 0.00 | 0.50 ± 0.84 | 0.42 ± 0.30 | 0.00 ± 0.00 | 0.13 ± 0.28 | 0.33 ± 0.41 | 0.00 ± 0.00 | 0.53 ± 1.00 | 0.07 ± 0.12 | 0.00 ± 0.00 | 0.05 ± 0.08 | 0.07 ± 0.08 | 0.00 ± 0.00 | 0.07 ± 0.12 | 0.28 ± 0.38 |
| *Scabiosa ochroleuca* | S | 0.15 ± 0.21 | 0.67 ± 0.76 | 0.38 ± 0.40 | 0.25 ± 0.61 | 0.98 ± 1.04 | 0.67 ± 0.53 | 1.92 ± 2.54 | 1.97 ± 2.35 | 1.42 ± 0.93 | 0.57 ± 0.78 | 0.83 ± 0.93 | 0.65 ± 0.51 | 2.42 ± 3.35 | 2.05 ± 2.84 | 1.27 ± 1.77 |
| *Securigera varia* | S | 0.50 ± 1.22 | 0.00 ± 0.00 | 0.05 ± 0.12 | 0.00 ± 0.00 | 0.00 ± 0.00 | 0.05 ± 0.12 | 12.17 ± 28.34 | 1.17 ± 1.29 | 1.02 ± 1.97 | 2.58 ± 3.32 | 4.67 ± 7.76 | 1.97 ± 3.96 | 14.17 ± 25.58 | 10.08 ± 22.05 | 8.00 ± 14.78 |
| *Silene viscosa* | S | 0.00 ± 0.00 | 0.00 ± 0.00 | 0.00 ± 0.00 | 0.00 ± 0.00 | 0.50 ± 0.63 | 0.63 ± 0.43 | 0.13 ± 0.22 | 0.52 ± 0.60 | 0.90 ± 0.57 | 0.42 ± 0.80 | 0.33 ± 0.52 | 0.78 ± 0.78 | 0.38 ± 0.80 | 0.75 ± 0.69 | 0.92 ± 0.68 |
| *Silene vulgaris* | S | 1.25 ± 1.00 | 1.92 ± 2.06 | 3.28 ± 2.40 | 3.12 ± 1.91 | 3.00 ± 1.26 | 2.20 ± 1.66 | 3.92 ± 2.69 | 5.83 ± 3.82 | 6.33 ± 6.11 | 1.2 ± 0.65 | 2.00 ± 2.53 | 1.67 ± 0.88 | 2.80 ± 3.67 | 1.17 ± 1.21 | 2.95 ± 3.57 |
| *Trifolium angulatum* | S | 0.00 ± 0.00 | 0.00 ± 0.00 | 0.00 ± 0.00 | 0.00 ± 0.00 | 0.25 ± 0.42 | 0.00 ± 0.00 | 0.18 ± 0.40 | 0.17 ± 0.41 | 0.00 ± 0.00 | 0.67 ± 1.63 | 0.00 ± 0.00 | 0.02 ± 0.04 | 0.00 ± 0.00 | 0.02 ± 0.04 | 0.00 ± 0.00 |
| *Trifolium campestre* | S | 0.13 ± 0.22 | 0.18 ± 0.40 | 0.10 ± 0.15 | 0.00 ± 0.00 | 0.12 ± 0.29 | 0.07 ± 0.12 | 0.00 ± 0.00 | 0.05 ± 0.12 | 0.13 ± 0.28 | 0.00 ± 0.00 | 0.20 ± 0.40 | 0.00 ± 0.00 | 0.42 ± 0.80 | 0.8 ± 0.63 | 0.52 ± 0.52 |
| *Trifolium retusum* | S | 0.00 ± 0.00 | 0.02 ± 0.04 | 0.00 ± 0.00 | 0.00 ± 0.00 | 0.50 ± 1.22 | 0.00 ± 0.00 | 0.00 ± 0.00 | 0.00 ± 0.00 | 0.00 ± 0.00 | 0.12 ± 0.29 | 0.00 ± 0.00 | 0.00 ± 0.00 | 0.00 ± 0.00 | 0.00 ± 0.00 | 0.03 ± 0.05 |
| *Trifolium striatum* | S | 0.00 ± 0.00 | 0.02 ± 0.04 | 0.07 ± 0.12 | 0.30 ± 0.60 | 0.17 ± 0.41 | 0.50 ± 0.84 | 0.33 ± 0.82 | 0.02 ± 0.04 | 0.17 ± 0.41 | 0.00 ± 0.00 | 0.00 ± 0.00 | 0.00 ± 0.00 | 0.10 ± 0.15 | 1.13 ± 2.39 | 1.00 ± 1.97 |
| *Achillea setacea* | | 0.00 ± 0.00 | 0.33 ± 0.82 | 0.00 ± 0.00 | 0.00 ± 0.00 | 0.00 ± 0.00 | 0.00 ± 0.00 | 0.00 ± 0.00 | 0.17 ± 0.41 | 0.00 ± 0.00 | 0.00 ± 0.00 | 0.00 ± 0.00 | 0.02 ± 0.04 | 0.00 ± 0.00 | 0.83 ± 2.04 | 0.17 ± 0.41 |
| *Alopecurus pratensis* | | 0.00 ± 0.00 | 0.00 ± 0.00 | 0.00 ± 0.00 | 0.00 ± 0.00 | 0.12 ± 0.29 | 0.08 ± 0.20 | 0.17 ± 0.41 | 0.00 ± 0.00 | 0.00 ± 0.00 | 0.00 ± 0.00 | 0.00 ± 0.00 | 0.00 ± 0.00 | 0.00 ± 0.00 | 0.00 ± 0.00 | 0.00 ± 0.00 |
| *Amaranthus retroflexus* | W | 0.08 ± 0.20 | 0.00 ± 0.00 | 0.02 ± 0.04 | 0.00 ± 0.00 | 0.00 ± 0.00 | 0.00 ± 0.00 | 0.00 ± 0.00 | 0.00 ± 0.00 | 0.00 ± 0.00 | 0.00 ± 0.00 | 0.00 ± 0.00 | 0.00 ± 0.00 | 0.00 ± 0.00 | 0.00 ± 0.00 | 0.00 ± 0.00 |
| *Anagallis arvensis* | W | 0.13 ± 0.22 | 0.2.0 ± 0.27 | 0.23 ± 0.27 | 0.00 ± 0.00 | 0.08 ± 0.20 | 0.00 ± 0.00 | 0.00 ± 0.00 | 0.00 ± 0.00 | 0.00 ± 0.00 | 0.00 ± 0.00 | 0.00 ± 0.00 | 0.00 ± 0.00 | 0.00 ± 0.00 | 0.00 ± 0.00 | 0.00 ± 0.00 |
| *Anthemis arvensis* | W | 0.00 ± 0.00 | 0.00 ± 0.00 | 0.00 ± 0.00 | 0.00 ± 0.00 | 0.12 ± 0.29 | 1.00 ± 2.45 | 0.00 ± 0.00 | 0.00 ± 0.00 | 0.00 ± 0.00 | 0.00 ± 0.00 | 0.00 ± 0.00 | 0.00 ± 0.00 | 0.00 ± 0.00 | 0.00 ± 0.00 | 0.00 ± 0.00 |
| *Anthriscus caucalis* | W | 0.00 ± 0.00 | 0.00 ± 0.00 | 0.00 ± 0.00 | 0.00 ± 0.00 | 0.02 ± 0.04 | 0.05 ± 0.12 | 0.00 ± 0.00 | 0.00 ± 0.00 | 0.00 ± 0.00 | 0.00 ± 0.00 | 0.00 ± 0.00 | 0.00 ± 0.00 | 0.00 ± 0.00 | 0.00 ± 0.00 | 0.00 ± 0.00 |
| *Arabidopsis thaliana* | | 0.00 ± 0.00 | 0.00 ± 0.00 | 0.00 ± 0.00 | 0.25 ± 0.61 | 0.27 ± 0.41 | 0.40 ± 0.45 | 0.00 ± 0.00 | 0.00 ± 0.00 | 0.00 ± 0.00 | 0.00 ± 0.00 | 0.00 ± 0.00 | 0.00 ± 0.00 | 0.00 ± 0.00 | 0.00 ± 0.00 | 0.00 ± 0.00 |
| *Arabis hirsuta* | | 0.00 ± 0.00 | 0.00 ± 0.00 | 0.00 ± 0.00 | 0.00 ± 0.00 | 0.00 ± 0.00 | 0.00 ± 0.00 | 0.00 ± 0.00 | 0.00 ± 0.00 | 0.00 ± 0.00 | 0.00 ± 0.00 | 0.00 ± 0.00 | 0.00 ± 0.00 | 0.00 ± 0.00 | 0.05 ± 0.12 | 0.00 ± 0.00 |
| *Arenaria serpyllifolia* | | 0.00 ± 0.00 | 0.00 ± 0.00 | 0.00 ± 0.00 | 0.40 ± 0.79 | 0.12 ± 0.29 | 0.33 ± 0.82 | 0.17 ± 0.29 | 0.02 ± 0.04 | 0.08 ± 0.20 | 0.00 ± 0.00 | 0.33 ± 0.61 | 0.12 ± 0.29 | 0.12 ± 0.29 | 0.02 ± 0.04 | 0.00 ± 0.00 |
| *Bilderdykia convolvulus* | W | 1.75 ± 3.56 | 0.58 ± 0.92 | 1.17 ± 2.40 | 0.05 ± 0.12 | 0.00 ± 0.00 | 0.00 ± 0.00 | 0.00 ± 0.00 | 0.00 ± 0.00 | 0.00 ± 0.00 | 0.00 ± 0.00 | 0.00 ± 0.00 | 0.00 ± 0.00 | 0.00 ± 0.00 | 0.00 ± 0.00 | 0.00 ± 0.00 |
| *Bromus arvensis* | W | 0.00 ± 0.00 | 0.00 ± 0.00 | 0.00 ± 0.00 | 0.00 ± 0.00 | 0.02 ± 0.04 | 0.15 ± 0.21 | 0.00 ± 0.00 | 0.00 ± 0.00 | 0.00 ± 0.00 | 0.00 ± 0.00 | 0.00 ± 0.00 | 0.00 ± 0.00 | 0.00 ± 0.00 | 0.00 ± 0.00 | 0.00 ± 0.00 |
| *Bromus inermis* | | 1.00 ± 1.55 | 0.30 ± 0.40 | 0.08 ± 0.20 | 3.22 ± 5.98 | 0.75 ± 1.17 | 0.42 ± 0.49 | 1.50 ± 2.81 | 0.83 ± 1.33 | 1.83 ± 1.94 | 1.88 ± 4.00 | 0.72 ± 1.18 | 0.95 ± 0.99 | 1.00 ± 2.00 | 1.27 ± 1.59 | 1.02 ± 1.57 |

| | | | | | | | | | | | | | | | | |
|---|---|---|---|---|---|---|---|---|---|---|---|---|---|---|---|---|
| *Bromus mollis* | | 0.00 ± 0.00 | 0.00 ± 0.00 | 0.02 ± 0.04 | 4.22 ± 10.18 | 0.63 ± 0.77 | 0.78 ± 1.13 | 2.05 ± 4.88 | 0.00 ± 0.00 | 0.37 ± 0.80 | 0.13 ± 0.22 | 0.17 ± 0.41 | 0.12 ± 0.19 | 0.00 ± 0.00 | 0.00 ± 0.00 | 0.08 ± 0.20 |
| *Bromus sterilis* | W | 0.00 ± 0.00 | 0.00 ± 0.00 | 0.00 ± 0.00 | 0.00 ± 0.00 | 0.12 ± 0.29 | 0.12 ± 0.29 | 1.67 ± 4.08 | 0.00 ± 0.00 | 0.00 ± 0.00 | 0.00 ± 0.00 | 0.00 ± 0.00 | 0.00 ± 0.00 | 0.00 ± 0.00 | 0.00 ± 0.00 | 0.00 ± 0.00 |
| *Bromus tectorum* | | 0.00 ± 0.00 | 0.00 ± 0.00 | 0.00 ± 0.00 | 3.17 ± 6.01 | 0.00 ± 0.00 | 0.88 ± 2.02 | 0.50 ± 1.22 | 0.00 ± 0.00 | 0.00 ± 0.00 | 0.00 ± 0.00 | 0.00 ± 0.00 | 0.00 ± 0.00 | 0.00 ± 0.00 | 0.00 ± 0.00 | 0.00 ± 0.00 |
| *Camelina microcarpa* | W | 0.00 ± 0.00 | 0.00 ± 0.00 | 0.00 ± 0.00 | 0.00 ± 0.00 | 0.17 ± 0.41 | 0.17 ± 0.41 | 0.00 ± 0.00 | 0.00 ± 0.00 | 0.00 ± 0.00 | 0.00 ± 0.00 | 0.00 ± 0.00 | 0.00 ± 0.00 | 0.00 ± 0.00 | 0.00 ± 0.00 | 0.00 ± 0.00 |
| *Capsella bursa pastoris* | W | 1.03 ± 2.43 | 0.10 ± 0.15 | 0.18 ± 0.18 | 2.75 ± 2.56 | 2.05 ± 2.30 | 4.92 ± 2.87 | 0.18 ± 0.40 | 0.00 ± 0.00 | 0.02 ± 0.04 | 0.00 ± 0.00 | 0.00 ± 0.00 | 0.00 ± 0.00 | 0.00 ± 0.00 | 0.00 ± 0.00 | 0.00 ± 0.00 |
| *Carduus acanthoides* | W | 0.55 ± 1.21 | 0.00 ± 0.00 | 0.00 ± 0.00 | 0.75 ± 1.25 | 0.00 ± 0.00 | 1.12 ± 1.61 | 0.00 ± 0.00 | 0.67 ± 1.63 | 0.10 ± 0.20 | 0.05 ± 0.12 | 0.10 ± 0.15 | 3.43 ± 8.12 | 0.00 ± 0.00 | 0.08 ± 0.20 | 0.23 ± 0.36 |
| *Cerastium dubium* | | 0.00 ± 0.00 | 0.00 ± 0.00 | 0.00 ± 0.00 | 0.00 ± 0.00 | 0.02 ± 0.04 | 0.00 ± 0.00 | 0.00 ± 0.00 | 0.00 ± 0.00 | 0.00 ± 0.00 | 0.00 ± 0.00 | 0.00 ± 0.00 | 0.00 ± 0.00 | 0.00 ± 0.00 | 0.00 ± 0.00 | 0.00 ± 0.00 |
| *Cerastium semidecandrum* | | 0.00 ± 0.00 | 0.00 ± 0.00 | 0.02 ± 0.04 | 0.83 ± 1.33 | 0.7 ± 1.17 | 0.35 ± 0.27 | 0.33 ± 0.82 | 0.00 ± 0.00 | 0.08 ± 0.20 | 0.00 ± 0.00 | 0.00 ± 0.00 | 0.00 ± 0.00 | 0.00 ± 0.00 | 0.00 ± 0.00 | 0.00 ± 0.00 |
| *Chenopodium album* | W | 0.13 ± 0.28 | 0.22 ± 0.25 | 0.33 ± 0.31 | 0.00 ± 0.00 | 0.00 ± 0.00 | 0.00 ± 0.00 | 0.00 ± 0.00 | 0.00 ± 0.00 | 0.00 ± 0.00 | 0.00 ± 0.00 | 0.00 ± 0.00 | 0.05 ± 0.08 | 0.00 ± 0.00 | 0.00 ± 0.00 | 0.00 ± 0.00 |
| *Chenopodium hybridum* | W | 0.08 ± 0.20 | 0.00 ± 0.00 | 0.08 ± 0.20 | 0.00 ± 0.00 | 0.00 ± 0.00 | 0.00 ± 0.00 | 0.00 ± 0.00 | 0.00 ± 0.00 | 0.00 ± 0.00 | 0.00 ± 0.00 | 0.00 ± 0.00 | 0.00 ± 0.00 | 0.00 ± 0.00 | 0.00 ± 0.00 | 0.00 ± 0.00 |
| *Cirsium arvense* | W | 3.42 ± 4.98 | 5.22 ± 12.14 | 8.60 ± 7.22 | 2.00 ± 3.95 | 0.62 ± 1.20 | 3.30 ± 3.90 | 0.22 ± 0.40 | 0.60 ± 1.19 | 0.35 ± 0.57 | 0.00 ± 0.00 | 0.13 ± 0.22 | 0.13 ± 0.19 | 0.00 ± 0.00 | 0.02 ± 0.04 | 0.03 ± 0.05 |
| *Cirsium vulgare* | W | 0.00 ± 0.00 | 0.00 ± 0.00 | 0.00 ± 0.00 | 0.00 ± 0.00 | 0.00 ± 0.00 | 0.50 ± 1.22 | 0.00 ± 0.00 | 0.00 ± 0.00 | 0.00 ± 0.00 | 0.03 ± 0.08 | 0.00 ± 0.00 | 0.00 ± 0.00 | 0.00 ± 0.00 | 0.00 ± 0.00 | 0.00 ± 0.00 |
| *Consolida regalis* | W | 0.00 ± 0.00 | 0.38 ± 0.8 | 0.20 ± 0.32 | 0.08 ± 0.20 | 0.02 ± 0.04 | 0.03 ± 0.05 | 0.00 ± 0.00 | 0.00 ± 0.00 | 0.00 ± 0.00 | 0.02 ± 0.04 | 0.08 ± 0.20 | 0.00 ± 0.00 | 0.00 ± 0.00 | 0.00 ± 0.00 | 0.00 ± 0.00 |
| *Convolvulus arvensis* | W | 1.95 ± 2.31 | 5.50 ± 8.98 | 3.17 ± 4.67 | 3.08 ± 4.63 | 3.67 ± 4.03 | 1.75 ± 1.78 | 1.00 ± 0.84 | 1.58 ± 2.22 | 0.38 ± 0.38 | 0.18 ± 0.21 | 1.3 ± 1.85 | 0.53 ± 1.21 | 0.38 ± 0.49 | 0.13 ± 0.28 | 0.25 ± 0.27 |
| *Conyza canadensis* | W | 0.12 ± 0.29 | 0.17 ± 0.41 | 0.13 ± 0.22 | 0.25 ± 0.61 | 0.00 ± 0.00 | 0.00 ± 0.00 | 0.10 ± 0.20 | 0.23 ± 0.39 | 0.07 ± 0.12 | 0.07 ± 0.12 | 1.08 ± 2.41 | 0.35 ± 0.81 | 0.00 ± 0.00 | 0.00 ± 0.00 | 0.00 ± 0.00 |
| *Crepis setosa* | W | 0.00 ± 0.00 | 0.00 ± 0.00 | 0.00 ± 0.00 | 0.00 ± 0.00 | 0.00 ± 0.00 | 0.00 ± 0.00 | 0.00 ± 0.00 | 0.67 ± 1.63 | 0.00 ± 0.00 | 0.00 ± 0.00 | 0.00 ± 0.00 | 0.00 ± 0.00 | 0.05 ± 0.12 | 0.25 ± 0.61 | 0.05 ± 0.12 |
| *Crepis tectorum* | W | 0.25 ± 0.61 | 2.00 ± 4.90 | 0.53 ± 0.64 | 0.00 ± 0.00 | 0.00 ± 0.00 | 0.00 ± 0.00 | 0.00 ± 0.00 | 0.00 ± 0.00 | 0.00 ± 0.00 | 0.00 ± 0.00 | 0.00 ± 0.00 | 0.00 ± 0.00 | 0.00 ± 0.00 | 0.00 ± 0.00 | 0.00 ± 0.00 |
| *Cruciata pedemontana* | | 0.05 ± 0.12 | 0.02 ± 0.04 | 0.00 ± 0.00 | 0.63 ± 0.89 | 0.33 ± 0.31 | 1.43 ± 3.22 | 0.08 ± 0.20 | 0.00 ± 0.00 | 0.00 ± 0.00 | 0.00 ± 0.00 | 0.00 ± 0.00 | 0.03 ± 0.08 | 0.00 ± 0.00 | 0.00 ± 0.00 | 0.00 ± 0.00 |
| *Cynodon dactylon* | W | 0.00 ± 0.00 | 0.08 ± 0.20 | 0.00 ± 0.00 | 0.00 ± 0.00 | 0.00 ± 0.00 | 0.08 ± 0.20 | 0.00 ± 0.00 | 0.00 ± 0.00 | 0.17 ± 0.41 | 0.00 ± 0.00 | 0.00 ± 0.00 | 0.00 ± 0.00 | 0.00 ± 0.00 | 0.00 ± 0.00 | 0.00 ± 0.00 |
| *Cynoglossum officinale* | W | 0.00 ± 0.00 | 2.33 ± 5.72 | 0.00 ± 0.00 | 0.00 ± 0.00 | 0.00 ± 0.00 | 2.75 ± 6.49 | 0.00 ± 0.00 | 0.58 ± 1.43 | 0.05 ± 0.12 | 0.05 ± 0.12 | 0.12 ± 0.20 | 0.05 ± 0.12 | 0.00 ± 0.00 | 0.08 ± 0.20 | 0.00 ± 0.00 |
| *Dactylis glomerata* | | 0.00 ± 0.00 | 0.00 ± 0.00 | 0.02 ± 0.04 | 0.00 ± 0.00 | 0.02 ± 0.04 | 0.13 ± 0.22 | 0.00 ± 0.00 | 0.50 ± 1.22 | 0.25 ± 0.61 | 0.07 ± 0.12 | 0.27 ± 0.61 | 0.07 ± 0.16 | 0.63 ± 0.89 | 0.50 ± 1.22 | 0.45 ± 0.81 |
| *Datura stramonium* | W | 0.00 ± 0.00 | 0.17 ± 0.41 | 0.17 ± 0.41 | 0.00 ± 0.00 | 0.00 ± 0.00 | 0.00 ± 0.00 | 0.00 ± 0.00 | 0.00 ± 0.00 | 0.00 ± 0.00 | 0.00 ± 0.00 | 0.00 ± 0.00 | 0.00 ± 0.00 | 0.00 ± 0.00 | 0.00 ± 0.00 | 0.00 ± 0.00 |
| *Daucus carota* | | 0.00 ± 0.00 | 1.02 ± 2.44 | 0.07 ± 0.12 | 0.00 ± 0.00 | 0.12 ± 0.29 | 0.00 ± 0.00 | 0.00 ± 0.00 | 0.33 ± 0.82 | 0.05 ± 0.12 | 0.00 ± 0.00 | 0.03 ± 0.08 | 0.05 ± 0.08 | 0.00 ± 0.00 | 0.05 ± 0.12 | 0.00 ± 0.00 |
| *Descurainia sophia* | W | 0.05 ± 0.12 | 0.08 ± 0.20 | 0.00 ± 0.00 | 0.02 ± 0.04 | 0.85 ± 2.03 | 0.17 ± 0.29 | 0.00 ± 0.00 | 0.00 ± 0.00 | 0.00 ± 0.00 | 0.00 ± 0.00 | 0.00 ± 0.00 | 0.00 ± 0.00 | 0.00 ± 0.00 | 0.00 ± 0.00 | 0.00 ± 0.00 |
| *Echinochloa crus gallii* | W | 0.20 ± 0.32 | 0.28 ± 0.34 | 0.20 ± 0.32 | 0.00 ± 0.00 | 0.00 ± 0.00 | 0.28 ± 0.45 | 0.00 ± 0.00 | 0.25 ± 0.61 | 0.05 ± 0.12 | 0.00 ± 0.00 | 0.00 ± 0.00 | 0.00 ± 0.00 | 0.08 ± 0.2 | 0.02 ± 0.04 | 0.02 ± 0.04 |
| *Elymus repens* | W | 0.33 ± 0.82 | 0.00 ± 0.00 | 0.12 ± 0.29 | 0.00 ± 0.00 | 0.38 ± 0.80 | 0.02 ± 0.04 | 0.00 ± 0.00 | 0.17 ± 0.41 | 0.08 ± 0.20 | 0.25 ± 0.61 | 0.00 ± 0.00 | 0.00 ± 0.00 | 0.00 ± 0.00 | 0.00 ± 0.00 | 0.02 ± 0.04 |
| *Epilobium tetragonum* | | 0.00 ± 0.00 | 0.02 ± 0.04 | 0.00 ± 0.00 | 0.00 ± 0.00 | 0.00 ± 0.00 | 0.05 ± 0.12 | 0.00 ± 0.00 | 0.00 ± 0.00 | 0.00 ± 0.00 | 0.00 ± 0.00 | 0.00 ± 0.00 | 0.00 ± 0.00 | 0.00 ± 0.00 | 0.00 ± 0.00 | 0.00 ± 0.00 |
| *Erophila verna* | | 0.00 ± 0.00 | 0.00 ± 0.00 | 0.00 ± 0.00 | 0.00 ± 0.00 | 0.00 ± 0.00 | 0.02 ± 0.04 | 0.00 ± 0.00 | 0.00 ± 0.00 | 0.00 ± 0.00 | 0.00 ± 0.00 | 0.00 ± 0.00 | 0.00 ± 0.00 | 0.00 ± 0.00 | 0.00 ± 0.00 | 0.00 ± 0.00 |
| *Eryngium campestre* | | 0.03 ± 0.05 | 0.00 ± 0.00 | 0.05 ± 0.05 | 0.00 ± 0.00 | 0.00 ± 0.00 | 0.02 ± 0.04 | 0.00 ± 0.00 | 0.00 ± 0.00 | 0.00 ± 0.00 | 0.00 ± 0.00 | 0.00 ± 0.00 | 0.00 ± 0.00 | 0.00 ± 0.00 | 0.00 ± 0.00 | 0.00 ± 0.00 |
| *Erysimum repandum* | W | 0.12 ± 0.29 | 0.08 ± 0.20 | 0.00 ± 0.00 | 0.00 ± 0.00 | 0.18 ± 0.4 | 0.02 ± 0.04 | 0.00 ± 0.00 | 0.00 ± 0.00 | 0.00 ± 0.00 | 0.00 ± 0.00 | 0.00 ± 0.00 | 0.00 ± 0.00 | 0.00 ± 0.00 | 0.00 ± 0.00 | 0.00 ± 0.00 |
| *Festuca arundinacea* | | 0.83 ± 1.33 | 0.00 ± 0.00 | 0.17 ± 0.41 | 0.08 ± 0.20 | 0.00 ± 0.00 | 0.00 ± 0.00 | 0.00 ± 0.00 | 0.00 ± 0.00 | 0.08 ± 0.20 | 0.05 ± 0.12 | 0.00 ± 0.00 | 0.00 ± 0.00 | 0.00 ± 0.00 | 0.00 ± 0.00 | 0.00 ± 0.00 |
| *Festuca pratensis* | | 0.00 ± 0.00 | 0.00 ± 0.00 | 0.00 ± 0.00 | 0.00 ± 0.00 | 0.00 ± 0.00 | 0.00 ± 0.00 | 0.00 ± 0.00 | 0.00 ± 0.00 | 0.00 ± 0.00 | 0.00 ± 0.00 | 0.00 ± 0.00 | 0.00 ± 0.00 | 0.67 ± 1.63 | 0.33 ± 0.82 | 0.00 ± 0.00 |
| *Festuca pseudovina* | | 0.08 ± 0.20 | 0.5 ± 0.84 | 1.00 ± 1.55 | 0.50 ± 1.22 | 0.58 ± 1.02 | 1.02 ± 2.44 | 1.25 ± 2.82 | 1.00 ± 2.00 | 0.50 ± 0.84 | 2.50 ± 4.81 | 5.83 ± 9.17 | 4.17 ± 6.65 | 1.83 ± 4.02 | 4.67 ± 7.66 | 1.67 ± 2.66 |
| *Festuca rupicola* | | 1.83 ± 4.02 | 1.28 ± 1.52 | 0.77 ± 0.97 | 4.55 ± 6.20 | 1.67 ± 1.86 | 2.17 ± 1.83 | 8.42 ± 9.41 | 3.25 ± 5.02 | 3.25 ± 3.66 | 36.33 ± 35.90 | 19.42 ± 32.17 | 20.83 ± 26.16 | 19.17 ± 21.54 | 15.83 ± 13.57 | 23.33 ± 26.77 |
| *Fumaria schleicheri* | | 0.00 ± 0.00 | 0.00 ± 0.00 | 0.00 ± 0.00 | 0.03 ± 0.05 | 0.22 ± 0.40 | 0.17 ± 0.41 | 0.00 ± 0.00 | 0.00 ± 0.00 | 0.00 ± 0.00 | 0.00 ± 0.00 | 0.00 ± 0.00 | 0.00 ± 0.00 | 0.00 ± 0.00 | 0.00 ± 0.00 | 0.00 ± 0.00 |
| *Galium aparine* | W | 0.00 ± 0.00 | 0.00 ± 0.00 | 0.00 ± 0.00 | 0.00 ± 0.00 | 0.00 ± 0.00 | 0.00 ± 0.00 | 0.02 ± 0.04 | 0.00 ± 0.00 | 0.00 ± 0.00 | 0.00 ± 0.00 | 0.00 ± 0.00 | 0.00 ± 0.00 | 0.00 ± 0.00 | 0.00 ± 0.00 | 0.00 ± 0.00 |
| *Galium spurium* | W | 0.02 ± 0.04 | 0.15 ± 0.21 | 0.02 ± 0.04 | 0.05 ± 0.12 | 0.05 ± 0.12 | 0.05 ± 0.12 | 0.00 ± 0.00 | 0.02 ± 0.04 | 0.02 ± 0.04 | 0.00 ± 0.00 | 0.08 ± 0.20 | 0.00 ± 0.00 | 0.00 ± 0.00 | 0.00 ± 0.00 | 0.00 ± 0.00 |
| *Geranium molle* | | 0.00 ± 0.00 | 0.00 ± 0.00 | 0.00 ± 0.00 | 0.02 ± 0.04 | 0.00 ± 0.00 | 0.00 ± 0.00 | 0.00 ± 0.00 | 0.00 ± 0.00 | 0.00 ± 0.00 | 0.00 ± 0.00 | 0.00 ± 0.00 | 0.00 ± 0.00 | 0.00 ± 0.00 | 0.00 ± 0.00 | 0.00 ± 0.00 |
| *Gypsophila muralis* | | 0.00 ± 0.00 | 0.17 ± 0.41 | 0.05 ± 0.12 | 0.00 ± 0.00 | 0.00 ± 0.00 | 0.00 ± 0.00 | 0.00 ± 0.00 | 0.02 ± 0.04 | 0.10 ± 0.20 | 0.00 ± 0.00 | 0.03 ± 0.08 | 0.05 ± 0.12 | 0.00 ± 0.00 | 0.00 ± 0.00 | 0.02 ± 0.04 |
| *Hibiscus trionum* | W | 0.13 ± 0.22 | 0.05 ± 0.12 | 0.15 ± 0.16 | 0.00 ± 0.00 | 0.00 ± 0.00 | 0.00 ± 0.00 | 0.00 ± 0.00 | 0.00 ± 0.00 | 0.00 ± 0.00 | 0.00 ± 0.00 | 0.00 ± 0.00 | 0.00 ± 0.00 | 0.00 ± 0.00 | 0.00 ± 0.00 | 0.00 ± 0.00 |
| *Hieracium echioides* | | 0.00 ± 0.00 | 0.00 ± 0.00 | 0.00 ± 0.00 | 0.00 ± 0.00 | 0.00 ± 0.00 | 0.00 ± 0.00 | 0.00 ± 0.00 | 0.00 ± 0.00 | 1.17 ± 2.86 | 0.00 ± 0.00 | 0.00 ± 0.00 | 0.00 ± 0.00 | 0.00 ± 0.00 | 0.00 ± 0.00 | 0.00 ± 0.00 |
| *Holosteum umbellatum* | W | 0.00 ± 0.00 | 0.00 ± 0.00 | 0.00 ± 0.00 | 0.12 ± 0.29 | 0.00 ± 0.00 | 0.00 ± 0.00 | 0.00 ± 0.00 | 0.00 ± 0.00 | 0.00 ± 0.00 | 0.00 ± 0.00 | 0.00 ± 0.00 | 0.00 ± 0.00 | 0.00 ± 0.00 | 0.00 ± 0.00 | 0.00 ± 0.00 |
| *Knautia arvensis* | | 0.00 ± 0.00 | 0.00 ± 0.00 | 0.00 ± 0.00 | 0.83 ± 0.94 | 0.00 ± 0.00 | 0.00 ± 0.00 | 2.58 ± 6.09 | 0.00 ± 0.00 | 0.00 ± 0.00 | 0.00 ± 0.00 | 0.00 ± 0.00 | 0.00 ± 0.00 | 0.00 ± 0.00 | 0.00 ± 0.00 | 0.00 ± 0.00 |
| *Koeleria cristata* | | 0.00 ± 0.00 | 0.00 ± 0.00 | 0.00 ± 0.00 | 0.00 ± 0.00 | 0.00 ± 0.00 | 0.00 ± 0.00 | 0.00 ± 0.00 | 0.00 ± 0.00 | 0.00 ± 0.00 | 0.02 ± 0.04 | 0.25 ± 0.61 | 0.00 ± 0.00 | 0.42 ± 0.66 | 0.67 ± 1.21 | 0.00 ± 0.00 |
| *Lactuca saligna* | | 0.00 ± 0.00 | 0.08 ± 0.20 | 0.23 ± 0.36 | 0.00 ± 0.00 | 0.00 ± 0.00 | 0.00 ± 0.00 | 0.00 ± 0.00 | 0.00 ± 0.00 | 0.25 ± 0.61 | 0.00 ± 0.00 | 0.00 ± 0.00 | 0.02 ± 0.04 | 0.00 ± 0.00 | 0.05 ± 0.12 | 0.00 ± 0.00 |
| *Lactuca serriola* | W | 0.08 ± 0.20 | 0.00 ± 0.00 | 0.22 ± 0.18 | 0.25 ± 0.61 | 0.05 ± 0.12 | 0.05 ± 0.12 | 0.00 ± 0.00 | 0.00 ± 0.00 | 0.00 ± 0.00 | 0.00 ± 0.00 | 0.02 ± 0.04 | 0.00 ± 0.00 | 0.00 ± 0.00 | 0.00 ± 0.00 | 0.00 ± 0.00 |
| *Lamium amplexicaule* | W | 0.00 ± 0.00 | 0.00 ± 0.00 | 0.00 ± 0.00 | 0.05 ± 0.12 | 0.33 ± 0.82 | 0.40 ± 0.35 | 0.00 ± 0.00 | 0.00 ± 0.00 | 0.00 ± 0.00 | 0.00 ± 0.00 | 0.00 ± 0.00 | 0.00 ± 0.00 | 0.00 ± 0.00 | 0.00 ± 0.00 | 0.00 ± 0.00 |
| *Lamium purpureum* | W | 0.00 ± 0.00 | 0.00 ± 0.00 | 0.00 ± 0.00 | 0.00 ± 0.00 | 0.00 ± 0.00 | 0.12 ± 0.29 | 0.00 ± 0.00 | 0.00 ± 0.00 | 0.00 ± 0.00 | 0.00 ± 0.00 | 0.00 ± 0.00 | 0.00 ± 0.00 | 0.00 ± 0.00 | 0.00 ± 0.00 | 0.00 ± 0.00 |
| *Lepidium draba* | W | 0.00 ± 0.00 | 0.00 ± 0.00 | 0.02 ± 0.04 | 0.00 ± 0.00 | 0.17 ± 0.41 | 0.15 ± 0.21 | 0.08 ± 0.20 | 0.00 ± 0.00 | 0.00 ± 0.00 | 0.42 ± 1.02 | 0.00 ± 0.00 | 0.00 ± 0.00 | 0.00 ± 0.00 | 0.00 ± 0.00 | 0.08 ± 0.20 |
| *Lepidium perfoliatum* | | 0.00 ± 0.00 | 0.25 ± 0.42 | 0.05 ± 0.12 | 0.00 ± 0.00 | 0.35 ± 0.81 | 0.05 ± 0.12 | 0.08 ± 0.20 | 0.00 ± 0.00 | 0.00 ± 0.00 | 0.50 ± 1.22 | 0.00 ± 0.00 | 0.00 ± 0.00 | 0.00 ± 0.00 | 0.00 ± 0.00 | 0.00 ± 0.00 |
| *Lepidium ruderale* | | 0.00 ± 0.00 | 0.00 ± 0.00 | 0.00 ± 0.00 | 0.00 ± 0.00 | 0.00 ± 0.00 | 0.00 ± 0.00 | 0.00 ± 0.00 | 0.00 ± 0.00 | 0.00 ± 0.00 | 0.00 ± 0.00 | 0.05 ± 0.12 | 0.05 ± 0.12 | 0.00 ± 0.00 | 0.00 ± 0.00 | 0.00 ± 0.00 |
| *Lithospermum arvense* | | 0.00 ± 0.00 | 0.00 ± 0.00 | 0.00 ± 0.00 | 0.00 ± 0.00 | 0.00 ± 0.00 | 0.02 ± 0.04 | 0.00 ± 0.00 | 0.00 ± 0.00 | 0.00 ± 0.00 | 0.00 ± 0.00 | 0.00 ± 0.00 | 0.00 ± 0.00 | 0.00 ± 0.00 | 0.00 ± 0.00 | 0.00 ± 0.00 |
| *Lycopsis arvensis* | W | 0.00 ± 0.00 | 0.00 ± 0.00 | 0.00 ± 0.00 | 0.00 ± 0.00 | 0.00 ± 0.00 | 0.00 ± 0.00 | 0.00 ± 0.00 | 0.05 ± 0.12 | 0.00 ± 0.00 | 0.00 ± 0.00 | 0.00 ± 0.00 | 0.00 ± 0.00 | 0.00 ± 0.00 | 0.00 ± 0.00 | 0.00 ± 0.00 |

| | | | | | | | | | | | | | | | | |
|---|---|---|---|---|---|---|---|---|---|---|---|---|---|---|---|---|
| *Matricaria chamomilla* | | 0.00 ± 0.00 | 0.00 ± 0.00 | 0.00 ± 0.00 | 0.00 ± 0.00 | 0.05 ± 0.12 | 0.00 ± 0.00 | 0.00 ± 0.00 | 0.02 ± 0.04 | 0.00 ± 0.00 | 0.00 ± 0.00 | 0.00 ± 0.00 | 0.00 ± 0.00 | 0.00 ± 0.00 | 0.00 ± 0.00 | 0.00 ± 0.00 |
| *Matricaria inodora* | W | 0.45 ± 0.81 | 1.63 ± 1.03 | 2.08 ± 3.01 | 0.10 ± 0.15 | 0.20 ± 0.20 | 1.83 ± 3.11 | 0.08 ± 0.20 | 0.83 ± 1.81 | 0.33 ± 0.52 | 0.00 ± 0.00 | 0.10 ± 0.20 | 0.07 ± 0.12 | 0.00 ± 0.00 | 0.00 ± 0.00 | 0.00 ± 0.00 |
| *Medicago falcata* | | 0.00 ± 0.00 | 0.00 ± 0.00 | 0.00 ± 0.00 | 0.00 ± 0.00 | 0.00 ± 0.00 | 0.00 ± 0.00 | 0.00 ± 0.00 | 0.00 ± 0.00 | 0.00 ± 0.00 | 0.08 ± 0.20 | 0.05 ± 0.12 | 0.00 ± 0.00 | 0.00 ± 0.00 | 0.00 ± 0.00 | 0.00 ± 0.00 |
| *Medicago lupulina* | | 0.08 ± 0.20 | 0.05 ± 0.12 | 0.02 ± 0.04 | 2.85 ± 6.00 | 0.30 ± 0.40 | 0.18 ± 0.40 | 0.33 ± 0.61 | 0.05 ± 0.12 | 0.05 ± 0.08 | 0.00 ± 0.00 | 0.00 ± 0.00 | 0.00 ± 0.00 | 0.05 ± 0.12 | 0.33 ± 0.61 | 0.13 ± 0.22 |
| *Medicago minima* | | 0.00 ± 0.00 | 0.00 ± 0.00 | 0.00 ± 0.00 | 0.00 ± 0.00 | 0.00 ± 0.00 | 0.00 ± 0.00 | 0.00 ± 0.00 | 0.00 ± 0.00 | 0.00 ± 0.00 | 0.17 ± 0.41 | 0.02 ± 0.04 | 0.03 ± 0.08 | 0.00 ± 0.00 | 0.00 ± 0.00 | 0.00 ± 0.00 |
| *Medicago sativa* | | 0.08 ± 0.20 | 0.08 ± 0.20 | 0.05 ± 0.12 | 0.17 ± 0.41 | 0.00 ± 0.00 | 0.08 ± 0.20 | 0.00 ± 0.00 | 0.00 ± 0.00 | 0.12 ± 0.29 | 0.00 ± 0.00 | 0.08 ± 0.20 | 0.08 ± 0.20 | 0.28 ± 0.45 | 0.05 ± 0.12 | 0.17 ± 0.26 |
| *Melandrium album* | W | 0.05 ± 0.12 | 0.05 ± 0.12 | 0.00 ± 0.00 | 0.08 ± 0.20 | 0.45 ± 0.81 | 0.17 ± 0.41 | 0.00 ± 0.00 | 0.08 ± 0.20 | 0.17 ± 0.41 | 0.00 ± 0.00 | 0.00 ± 0.00 | 0.00 ± 0.00 | 0.00 ± 0.00 | 0.00 ± 0.00 | 0.00 ± 0.00 |
| *Melilotus officinalis* | W | 0.00 ± 0.00 | 0.00 ± 0.00 | 0.00 ± 0.00 | 0.00 ± 0.00 | 0.00 ± 0.00 | 0.00 ± 0.00 | 0.00 ± 0.00 | 0.00 ± 0.00 | 0.00 ± 0.00 | 0.00 ± 0.00 | 0.00 ± 0.00 | 0.02 ± 0.04 | 0.00 ± 0.00 | 0.00 ± 0.00 | 0.00 ± 0.00 |
| *Myosotis stricta* | | 0.00 ± 0.00 | 0.00 ± 0.00 | 0.00 ± 0.00 | 0.67 ± 1.63 | 0.02 ± 0.04 | 0.18 ± 0.28 | 0.00 ± 0.00 | 0.00 ± 0.00 | 0.00 ± 0.00 | 0.00 ± 0.00 | 0.00 ± 0.00 | 0.00 ± 0.00 | 0.00 ± 0.00 | 0.00 ± 0.00 | 0.00 ± 0.00 |
| *Nigella sativa* | W | 0.00 ± 0.00 | 0.00 ± 0.00 | 0.00 ± 0.00 | 0.00 ± 0.00 | 0.00 ± 0.00 | 0.00 ± 0.00 | 0.00 ± 0.00 | 0.00 ± 0.00 | 0.00 ± 0.00 | 0.02 ± 0.04 | 0.00 ± 0.00 | 0.00 ± 0.00 | 0.00 ± 0.00 | 0.00 ± 0.00 | 0.00 ± 0.00 |
| *Papaver rhoeas* | W | 0.67 ± 1.63 | 0.25 ± 0.61 | 0.00 ± 0.00 | 0.00 ± 0.00 | 0.07 ± 0.12 | 0.05 ± 0.12 | 0.00 ± 0.00 | 0.00 ± 0.00 | 0.00 ± 0.00 | 0.00 ± 0.00 | 0.00 ± 0.00 | 0.00 ± 0.00 | 0.00 ± 0.00 | 0.00 ± 0.00 | 0.00 ± 0.00 |
| *Phragmites communis* | | 1.08 ± 2.01 | 0.00 ± 0.00 | 0.00 ± 0.00 | 0.42 ± 0.80 | 0.00 ± 0.00 | 0.00 ± 0.00 | 0.17 ± 0.41 | 0.12 ± 0.29 | 0.08 ± 0.20 | 0.87 ± 2.03 | 0.12 ± 0.29 | 0.00 ± 0.00 | 1.67 ± 4.08 | 0.05 ± 0.12 | 0.00 ± 0.00 |
| *Picris hieracioides* | | 0.12 ± 0.29 | 0.67 ± 1.21 | 0.82 ± 1.12 | 7.5 ± 11.73 | 0.83 ± 1.33 | 1.13 ± 1.47 | 4.5 ± 8.09 | 4.58 ± 7.67 | 0.83 ± 0.68 | 3.47 ± 7.63 | 2.23 ± 5.28 | 1.85 ± 4.48 | 1.00 ± 2.45 | 0.90 ± 1.56 | 0.22 ± 0.31 |
| *Pimpinella saxifraga* | | 0.00 ± 0.00 | 0.00 ± 0.00 | 0.00 ± 0.00 | 0.00 ± 0.00 | 0.00 ± 0.00 | 0.00 ± 0.00 | 0.00 ± 0.00 | 0.00 ± 0.00 | 0.02 ± 0.04 | 0.00 ± 0.00 | 0.00 ± 0.00 | 0.00 ± 0.00 | 0.00 ± 0.00 | 0.00 ± 0.00 | 0.00 ± 0.00 |
| *Plantago lanceolata* | | 0.33 ± 0.52 | 1.00 ± 2.00 | 0.10 ± 0.15 | 0.67 ± 1.08 | 1.45 ± 3.22 | 0.17 ± 0.41 | 0.25 ± 0.61 | 0.58 ± 0.92 | 0.55 ± 1.21 | 0.25 ± 0.61 | 0.25 ± 0.42 | 0.10 ± 0.20 | 1.33 ± 3.27 | 0.67 ± 1.21 | 0.00 ± 0.00 |
| *Poa angustifolia* | | 0.45 ± 0.81 | 1.00 ± 1.98 | 0.25 ± 0.42 | 1.07 ± 2.42 | 2.17 ± 1.60 | 3.50 ± 2.74 | 2.33 ± 4.08 | 3.58 ± 1.96 | 1.83 ± 1.17 | 3.33 ± 3.39 | 3.83 ± 3.60 | 1.03 ± 1.65 | 11.33 ± 19.24 | 9.67 ± 9.07 | 5.50 ± 3.89 |
| *Poa pratensis* | | 0.12 ± 0.29 | 0.05 ± 0.12 | 0.83 ± 1.33 | 0.00 ± 0.00 | 0.00 ± 0.00 | 0.00 ± 0.00 | 0.00 ± 0.00 | 0.00 ± 0.00 | 0.00 ± 0.00 | 0.00 ± 0.00 | 0.00 ± 0.00 | 0.00 ± 0.00 | 0.00 ± 0.00 | 0.00 ± 0.00 | 0.00 ± 0.00 |
| *Polygonum aviculare* | W | 4.33 ± 5.75 | 2.33 ± 2.73 | 5.85 ± 6.35 | 0.05 ± 0.12 | 0.03 ± 0.05 | 0.00 ± 0.00 | 0.58 ± 1.43 | 0.08 ± 0.20 | 0.00 ± 0.00 | 0.83 ± 2.04 | 0.00 ± 0.00 | 0.00 ± 0.00 | 0.42 ± 1.02 | 0.00 ± 0.00 | 0.00 ± 0.00 |
| *Potentilla reptans* | | 0.00 ± 0.00 | 0.00 ± 0.00 | 0.00 ± 0.00 | 0.00 ± 0.00 | 0.00 ± 0.00 | 0.00 ± 0.00 | 0.00 ± 0.00 | 0.25 ± 0.61 | 0.05 ± 0.12 | 0.00 ± 0.00 | 0.00 ± 0.00 | 0.00 ± 0.00 | 0.00 ± 0.00 | 0.00 ± 0.00 | 0.00 ± 0.00 |
| *Rumex acetosa* | | 0.00 ± 0.00 | 0.00 ± 0.00 | 0.25 ± 0.42 | 0.00 ± 0.00 | 0.00 ± 0.00 | 0.00 ± 0.00 | 0.00 ± 0.00 | 0.00 ± 0.00 | 0.05 ± 0.12 | 0.00 ± 0.00 | 0.00 ± 0.00 | 0.00 ± 0.00 | 0.00 ± 0.00 | 0.00 ± 0.00 | 0.00 ± 0.00 |
| *Rumex crispus* | W | 0.00 ± 0.00 | 0.00 ± 0.00 | 0.00 ± 0.00 | 0.12 ± 0.29 | 0.00 ± 0.00 | 0.17 ± 0.29 | 0.00 ± 0.00 | 0.00 ± 0.00 | 0.03 ± 0.08 | 0.00 ± 0.00 | 0.00 ± 0.00 | 0.02 ± 0.04 | 0.08 ± 0.20 | 0.00 ± 0.00 | 0.00 ± 0.00 |
| *Salvia austriaca* | | 0.00 ± 0.00 | 0.00 ± 0.00 | 0.00 ± 0.00 | 0.00 ± 0.00 | 0.00 ± 0.00 | 0.00 ± 0.00 | 0.00 ± 0.00 | 0.00 ± 0.00 | 0.05 ± 0.12 | 0.00 ± 0.00 | 0.00 ± 0.00 | 0.00 ± 0.00 | 0.00 ± 0.00 | 0.00 ± 0.00 | 0.00 ± 0.00 |
| *Setaria glauca* | W | 3.83 ± 8.01 | 1.50 ± 1.00 | 1.83 ± 1.83 | 0.00 ± 0.00 | 0.00 ± 0.00 | 0.00 ± 0.00 | 0.00 ± 0.00 | 0.00 ± 0.00 | 0.25 ± 0.61 | 0.33 ± 0.82 | 0.25 ± 0.61 | 0.02 ± 0.04 | 0.08 ± 0.20 | 0.08 ± 0.20 | 0.03 ± 0.05 |
| *Setaria viridis* | W | 0.5 ± 0.84 | 0.83 ± 1.60 | 0.40 ± 0.45 | 0.00 ± 0.00 | 0.00 ± 0.00 | 0.00 ± 0.00 | 0.77 ± 1.83 | 1.03 ± 1.11 | 0.1 ± 0.20 | 0.00 ± 0.00 | 0.00 ± 0.00 | 0.00 ± 0.00 | 0.00 ± 0.00 | 0.00 ± 0.00 | 0.00 ± 0.00 |
| *Solanum nigrum* | W | 0.13 ± 0.28 | 0.05 ± 0.12 | 0.02 ± 0.04 | 0.00 ± 0.00 | 0.00 ± 0.00 | 0.00 ± 0.00 | 0.00 ± 0.00 | 0.00 ± 0.00 | 0.00 ± 0.00 | 0.00 ± 0.00 | 0.00 ± 0.00 | 0.00 ± 0.00 | 0.00 ± 0.00 | 0.00 ± 0.00 | 0.00 ± 0.00 |
| *Sonchus asper* | W | 0.00 ± 0.00 | 0.33 ± 0.82 | 0.00 ± 0.00 | 0.00 ± 0.00 | 0.00 ± 0.00 | 0.00 ± 0.00 | 0.00 ± 0.00 | 0.00 ± 0.00 | 0.00 ± 0.00 | 0.00 ± 0.00 | 0.00 ± 0.00 | 0.00 ± 0.00 | 0.00 ± 0.00 | 0.00 ± 0.00 | 0.00 ± 0.00 |
| *Sonchus oleraceus* | W | 0.00 ± 0.00 | 0.00 ± 0.00 | 0.00 ± 0.00 | 0.00 ± 0.00 | 0.00 ± 0.00 | 0.07 ± 0.12 | 0.05 ± 0.12 | 0.00 ± 0.00 | 0.00 ± 0.00 | 0.00 ± 0.00 | 0.00 ± 0.00 | 0.00 ± 0.00 | 0.00 ± 0.00 | 0.00 ± 0.00 | 0.00 ± 0.00 |
| *Spergularia rubra* | | 0.00 ± 0.00 | 0.02 ± 0.04 | 0.00 ± 0.00 | 0.00 ± 0.00 | 0.00 ± 0.00 | 0.00 ± 0.00 | 0.00 ± 0.00 | 0.00 ± 0.00 | 0.00 ± 0.00 | 0.00 ± 0.00 | 0.00 ± 0.00 | 0.00 ± 0.00 | 0.00 ± 0.00 | 0.00 ± 0.00 | 0.00 ± 0.00 |
| *Stachys annua* | W | 2.17 ± 5.31 | 0.92 ± 1.20 | 0.17 ± 0.41 | 0.00 ± 0.00 | 0.00 ± 0.00 | 0.00 ± 0.00 | 0.00 ± 0.00 | 0.00 ± 0.00 | 0.00 ± 0.00 | 0.00 ± 0.00 | 0.00 ± 0.00 | 0.00 ± 0.00 | 0.00 ± 0.00 | 0.00 ± 0.00 | 0.00 ± 0.00 |
| *Stellaria media* | | 0.00 ± 0.00 | 0.00 ± 0.00 | 0.00 ± 0.00 | 0.05 ± 0.12 | 1.78 ± 4.04 | 0.00 ± 0.00 | 0.00 ± 0.00 | 0.00 ± 0.00 | 0.00 ± 0.00 | 0.00 ± 0.00 | 0.00 ± 0.00 | 0.00 ± 0.00 | 0.00 ± 0.00 | 0.00 ± 0.00 | 0.00 ± 0.00 |
| *Stenactis annua* | W | 0.00 ± 0.00 | 0.00 ± 0.00 | 0.00 ± 0.00 | 0.00 ± 0.00 | 0.00 ± 0.00 | 0.00 ± 0.00 | 0.00 ± 0.00 | 0.33 ± 0.82 | 0.50 ± 1.22 | 0.00 ± 0.00 | 0.08 ± 0.20 | 1.50 ± 3.67 | 0.00 ± 0.00 | 0.02 ± 0.04 | 0.63 ± 1.41 |
| *Taraxacum officinale* | W | 0.05 ± 0.12 | 0.63 ± 1.19 | 0.12 ± 0.29 | 0.50 ± 1.22 | 4.42 ± 7.76 | 3.38 ± 5.88 | 0.42 ± 0.80 | 0.37 ± 0.62 | 0.25 ± 0.30 | 0.05 ± 0.12 | 0.13 ± 0.22 | 0.03 ± 0.05 | 1.03 ± 1.37 | 0.05 ± 0.12 | 0.07 ± 0.12 |
| *Thymus glabrescens* | | 0.00 ± 0.00 | 0.00 ± 0.00 | 0.02 ± 0.04 | 0.00 ± 0.00 | 0.00 ± 0.00 | 0.05 ± 0.12 | 0.00 ± 0.00 | 0.00 ± 0.00 | 0.00 ± 0.00 | 0.00 ± 0.00 | 0.00 ± 0.00 | 0.00 ± 0.00 | 0.00 ± 0.00 | 0.00 ± 0.00 | 0.00 ± 0.00 |
| *Thlaspi arvense* | W | 0.05 ± 0.12 | 0.07 ± 0.12 | 0.05 ± 0.12 | 0.08 ± 0.20 | 0.12 ± 0.29 | 1.22 ± 2.84 | 0.00 ± 0.00 | 0.00 ± 0.00 | 0.00 ± 0.00 | 0.00 ± 0.00 | 0.00 ± 0.00 | 0.00 ± 0.00 | 0.00 ± 0.00 | 0.00 ± 0.00 | 0.00 ± 0.00 |
| *Torilis arvensis* | W | 0.00 ± 0.00 | 0.00 ± 0.00 | 0.00 ± 0.00 | 0.00 ± 0.00 | 0.02 ± 0.04 | 0.02 ± 0.04 | 0.00 ± 0.00 | 0.00 ± 0.00 | 0.00 ± 0.00 | 0.00 ± 0.00 | 0.00 ± 0.00 | 0.00 ± 0.00 | 0.00 ± 0.00 | 0.00 ± 0.00 | 0.00 ± 0.00 |
| *Tragopogon dubius* | | 0.00 ± 0.00 | 0.00 ± 0.00 | 0.00 ± 0.00 | 0.00 ± 0.00 | 0.02 ± 0.04 | 0.00 ± 0.00 | 0.00 ± 0.00 | 0.00 ± 0.00 | 0.00 ± 0.00 | 0.00 ± 0.00 | 0.03 ± 0.08 | 0.08 ± 0.13 | 0.00 ± 0.00 | 0.00 ± 0.00 | 0.00 ± 0.00 |
| *Trifolium angustifolia* | | 0.00 ± 0.00 | 0.00 ± 0.00 | 0.00 ± 0.00 | 0.00 ± 0.00 | 0.00 ± 0.00 | 0.00 ± 0.00 | 0.00 ± 0.00 | 0.00 ± 0.00 | 0.02 ± 0.04 | 0.00 ± 0.00 | 0.00 ± 0.00 | 0.00 ± 0.00 | 0.00 ± 0.00 | 0.00 ± 0.00 | 0.00 ± 0.00 |
| *Trifolium arvense* | | 0.00 ± 0.00 | 0.05 ± 0.12 | 0.00 ± 0.00 | 0.00 ± 0.00 | 0.00 ± 0.00 | 0.00 ± 0.00 | 0.05 ± 0.12 | 0.00 ± 0.00 | 0.05 ± 0.12 | 0.08 ± 0.13 | 0.07 ± 0.12 | 0.05 ± 0.12 | 1.50 ± 3.21 | 0.83 ± 0.98 | 0.50 ± 0.84 |
| *Trifolium pratense* | | 0.00 ± 0.00 | 0.00 ± 0.00 | 0.00 ± 0.00 | 0.00 ± 0.00 | 0.00 ± 0.00 | 0.00 ± 0.00 | 0.00 ± 0.00 | 0.00 ± 0.00 | 0.00 ± 0.00 | 0.00 ± 0.00 | 0.00 ± 0.00 | 0.02 ± 0.04 | 0.00 ± 0.00 | 0.00 ± 0.00 | 0.05 ± 0.12 |
| *Trifolium repens* | | 0.00 ± 0.00 | 0.33 ± 0.82 | 0.00 ± 0.00 | 0.00 ± 0.00 | 3.00 ± 7.35 | 0.00 ± 0.00 | 0.00 ± 0.00 | 0.00 ± 0.00 | 0.02 ± 0.04 | 0.00 ± 0.00 | 0.00 ± 0.00 | 0.00 ± 0.00 | 0.50 ± 1.22 | 0.50 ± 0.63 | 0.42 ± 0.80 |
| *Trifolium strictum* | | 0.00 ± 0.00 | 0.00 ± 0.00 | 0.00 ± 0.00 | 0.00 ± 0.00 | 0.00 ± 0.00 | 0.05 ± 0.12 | 0.00 ± 0.00 | 0.00 ± 0.00 | 0.00 ± 0.00 | 0.00 ± 0.00 | 0.00 ± 0.00 | 0.00 ± 0.00 | 0.00 ± 0.00 | 0.00 ± 0.00 | 0.00 ± 0.00 |
| *Veronica arvensis* | | 0.00 ± 0.00 | 0.00 ± 0.00 | 0.02 ± 0.04 | 0.58 ± 0.74 | 0.75 ± 0.64 | 0.47 ± 0.37 | 0.33 ± 0.82 | 0.00 ± 0.00 | 0.05 ± 0.12 | 0.00 ± 0.00 | 0.00 ± 0.00 | 0.00 ± 0.00 | 0.00 ± 0.00 | 0.00 ± 0.00 | 0.00 ± 0.00 |
| *Veronica hederifolia* | W | 0.00 ± 0.00 | 0.00 ± 0.00 | 0.00 ± 0.00 | 0.12 ± 0.29 | 0.17 ± 0.41 | 0.23 ± 0.39 | 0.00 ± 0.00 | 0.00 ± 0.00 | 0.00 ± 0.00 | 0.00 ± 0.00 | 0.00 ± 0.00 | 0.00 ± 0.00 | 0.00 ± 0.00 | 0.00 ± 0.00 | 0.00 ± 0.00 |
| *Veronica polita* | W | 0.00 ± 0.00 | 0.00 ± 0.00 | 0.00 ± 0.00 | 0.00 ± 0.00 | 0.00 ± 0.00 | 0.02 ± 0.04 | 0.00 ± 0.00 | 0.00 ± 0.00 | 0.00 ± 0.00 | 0.00 ± 0.00 | 0.00 ± 0.00 | 0.00 ± 0.00 | 0.00 ± 0.00 | 0.00 ± 0.00 | 0.00 ± 0.00 |
| *Vicia angustifolia* | | 0.02 ± 0.04 | 0.00 ± 0.00 | 0.02 ± 0.04 | 0.08 ± 0.20 | 0.32 ± 0.37 | 0.30 ± 0.28 | 0.05 ± 0.12 | 0.47 ± 0.78 | 0.18 ± 0.28 | 0.00 ± 0.00 | 0.00 ± 0.00 | 0.00 ± 0.00 | 0.00 ± 0.00 | 0.50 ± 1.22 | 0.33 ± 0.82 |
| *Vicia grandiflora* | | 0.00 ± 0.00 | 0.00 ± 0.00 | 0.00 ± 0.00 | 0.00 ± 0.00 | 0.05 ± 0.12 | 0.28 ± 0.45 | 0.00 ± 0.00 | 0.00 ± 0.00 | 0.00 ± 0.00 | 0.00 ± 0.00 | 0.00 ± 0.00 | 0.00 ± 0.00 | 0.00 ± 0.00 | 0.00 ± 0.00 | 0.28 ± 0.45 |
| *Vicia hirsuta* | | 0.00 ± 0.00 | 0.07 ± 0.12 | 0.02 ± 0.04 | 0.92 ± 0.92 | 0.57 ± 0.58 | 0.97 ± 1.50 | 0.25 ± 0.61 | 0.40 ± 0.79 | 0.42 ± 1.02 | 0.00 ± 0.00 | 0.00 ± 0.00 | 0.02 ± 0.04 | 0.00 ± 0.00 | 0.17 ± 0.41 | 0.10 ± 0.20 |
| *Vicia lathyroides* | | 0.25 ± 0.61 | 0.12 ± 0.15 | 0.18 ± 0.28 | 0.50 ± 0.77 | 0.30 ± 0.40 | 0.20 ± 0.27 | 0.00 ± 0.00 | 0.00 ± 0.00 | 0.00 ± 0.00 | 0.00 ± 0.00 | 0.00 ± 0.00 | 0.00 ± 0.00 | 0.00 ± 0.00 | 0.00 ± 0.00 | 0.00 ± 0.00 |
| *Vicia pannonica* | W | 0.00 ± 0.00 | 0.00 ± 0.00 | 0.00 ± 0.00 | 0.00 ± 0.00 | 0.12 ± 0.29 | 0.08 ± 0.20 | 0.00 ± 0.00 | 0.00 ± 0.00 | 0.00 ± 0.00 | 0.00 ± 0.00 | 0.00 ± 0.00 | 0.00 ± 0.00 | 0.00 ± 0.00 | 0.00 ± 0.00 | 0.00 ± 0.00 |
| *Vicia tetrasperma* | | 0.00 ± 0.00 | 0.02 ± 0.04 | 0.00 ± 0.00 | 0.00 ± 0.00 | 0.13 ± 0.22 | 0.15 ± 0.21 | 0.00 ± 0.00 | 0.00 ± 0.00 | 0.02 ± 0.04 | 0.00 ± 0.00 | 0.00 ± 0.00 | 0.00 ± 0.00 | 0.00 ± 0.00 | 0.00 ± 0.00 | 0.02 ± 0.04 |
| *Vicia villosa* | W | 0.00 ± 0.00 | 0.00 ± 0.00 | 0.00 ± 0.00 | 0.42 ± 1.02 | 0.05 ± 0.12 | 0.05 ± 0.12 | 0.17 ± 0.41 | 0.17 ± 0.41 | 3.33 ± 8.16 | 0.00 ± 0.00 | 0.00 ± 0.00 | 0.00 ± 0.00 | 0.00 ± 0.00 | 0.00 ± 0.00 | 0.17 ± 0.26 |
| *Viola arvensis* | W | 0.00 ± 0.00 | 0.02 ± 0.04 | 0.00 ± 0.00 | 0.00 ± 0.00 | 0.00 ± 0.00 | 0.00 ± 0.00 | 0.00 ± 0.00 | 0.00 ± 0.00 | 0.00 ± 0.00 | 0.00 ± 0.00 | 0.00 ± 0.00 | 0.00 ± 0.00 | 0.00 ± 0.00 | 0.00 ± 0.00 | 0.00 ± 0.00 |
| *Viola kitaibeliana* | | 0.00 ± 0.00 | 0.00 ± 0.00 | 0.00 ± 0.00 | 0.00 ± 0.00 | 0.17 ± 0.41 | 0.00 ± 0.00 | 0.00 ± 0.00 | 0.00 ± 0.00 | 0.00 ± 0.00 | 0.00 ± 0.00 | 0.00 ± 0.00 | 0.00 ± 0.00 | 0.00 ± 0.00 | 0.00 ± 0.00 | 0.00 ± 0.00 |

**Table S2**. Effects of gap size and year on dependent variables related to the establishment success of the sown species, matrix grasses and weeds (Generalized Linear Mixed Models). Significant effects are marked with boldface. The arrows indicate the direction of vegetation changes with increasing gap size (size) or time (year) since the installation of the gaps.

| | Gap size | | Year | |
|---|---|---|---|---|
| | F | p | F | p |
| Total cover | 1.822 | 0.169 | **8.649** | **<0.001**↑ |
| Sown species cover | **5.441** | **0.006**↑ | **7.971** | **<0.001**↑ |
| Perennial sown | 2.644 | 0.078 | **14.587** | **<0.001**↑ |
| Short-lived sown | 0.279 | 0.757 | **5.499** | **0.001**↓ |
| Matrix grasses | 0.694 | 0.503 | **9.870** | **<0.001**↑ |
| Weed species cover | 0.157 | 0.855 | **9.189** | **<0.001**↓ |
| Perennial weeds | 0.003 | 0.997 | 1.561 | 0.194 |
| Short-lived weeds | 0.294 | 0.746 | **7.773** | **<0.001**↓ |

**Table S3**. Effects of distance from the gap, gap size and year on the dependent variables related to the colonization success (Generalized Linear Mixed Models). Significant effects are marked with boldface. The arrows indicate the direction of changes (↑ increase or ↓ decrease) with increasing distance from gaps or increasing gap size.

| | Distance from gap | | Gap size | | Year | |
|---|---|---|---|---|---|---|
| | F | p | F | p | F | p |
| Species number | **366.593** | **<0.001↓** | **44.342** | **<0.001↑** | **9.414** | **<0.001** |
| Individual number | **3052.579** | **<0.001↓** | **360.887** | **<0.001↑** | **340.97** | **<0.001** |
| Number of flowering species | **176.935** | **<0.001↓** | **29.214** | **<0.001↑** | **16.871** | **<0.001** |
| Flowering shoots | **3649.264** | **<0.001↓** | **324.912** | **<0.001↑** | **684.397** | **<0.001** |

**Table S4.** Effects of distance from the gap, gap size and year on the individual number of the nine most abundant species in the colonization plots (Generalised Linear Mixed Models). The arrows indicate the direction of changes (↑ increase or ↓ decrease) with increasing distance from gaps or increasing gap size.

| | Distance from gap | | Gap size | | Year | |
|---|---|---|---|---|---|---|
| | F | p | F | p | F | p |
| *Achillea collina* | 1110.74 | <0.001↓ | 345.752 | <0.001↑ | 13.058 | <0.001 |
| *Centaurea solstitialis* | 3478.433 | <0.001↓ | 33.517 | <0.001↑ | 519.023 | <0.001 |
| *Dianthus pontederae* | 6.759 | 0.009↓ | 7.551 | 0.001↑ | 86.977 | <0.001 |
| *Galium verum* | 506.627 | <0.001↓ | 78.944 | <0.001↓ | 64.192 | <0.001 |
| *Lathyrus tuberosus* | 6.600 | 0.010↓ | 159.121 | <0.001↓ | 7.837 | <0.001 |
| *Podospermum canum* | 12.319 | <0.001↓ | 24.172 | <0.001↑ | 144.602 | <0.001 |
| *Potentilla argentea* | 22.645 | <0.001↓ | 1.762 | 0.172 | 9.368 | <0.001 |
| *Trifolium campestre* | 158.7 | <0.001↓ | 229.403 | <0.001↑ | 123.918 | <0.001 |
| *Trifolium striatum* | 16.078 | <0.001↓ | 95.925 | <0.001↑ | 229.679 | <0.001 |

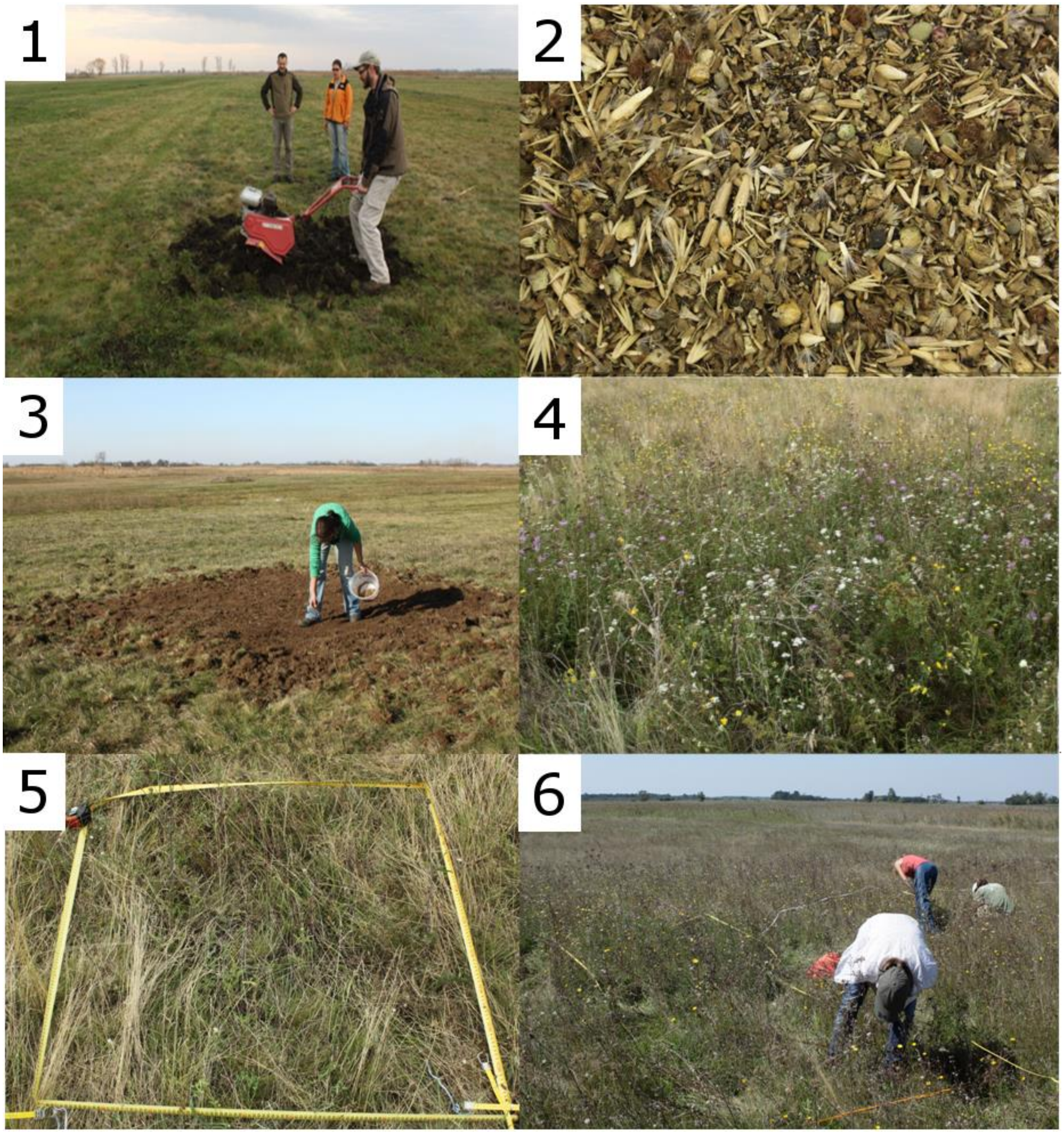


**Figure S1.** Moments of establishment gaps creation: 1 - Soil preparation, 2 - High-diversity seed mixture, 3 - Sowing the high diversity seed mixture, 4 - Establishment gap (16 $m^2$) in a species-poor grassland, 5 - Establishment gap (1 $m^2$) in a species-poor grassland, 6 - Monitoring the dispersal success of sown species.

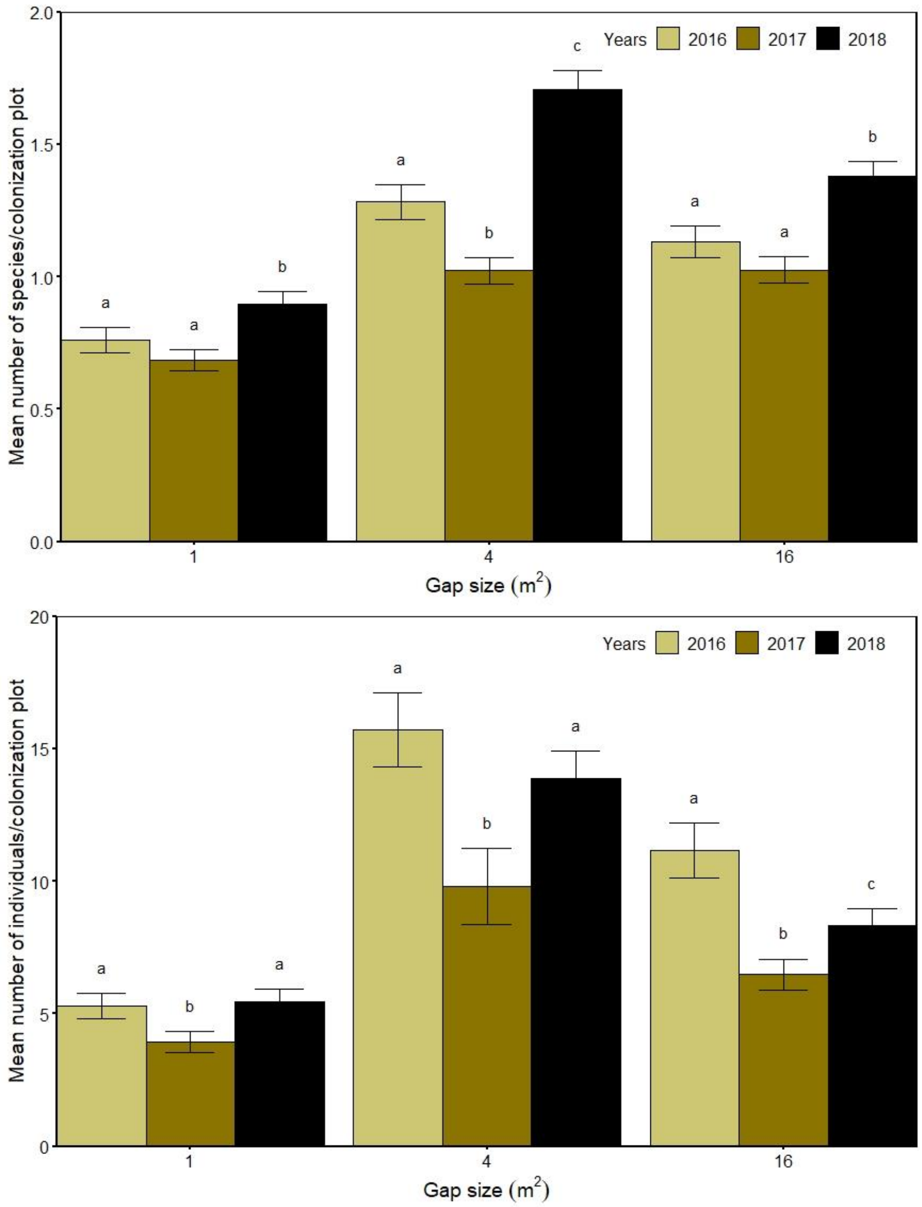


**Figure S2.** Number of sown species (mean ± SE) and number of individuals (mean ± SE) in the 1 m$^2$-sized colonization plots adjacent to different sized establishment gaps (1 m$^2$, 4 m$^2$ and 16 m$^2$) between 2016 and 2018. Letters indicate significant differences between the years within different gap sizes (GLMM and LSD-test, $p < 0.05$).

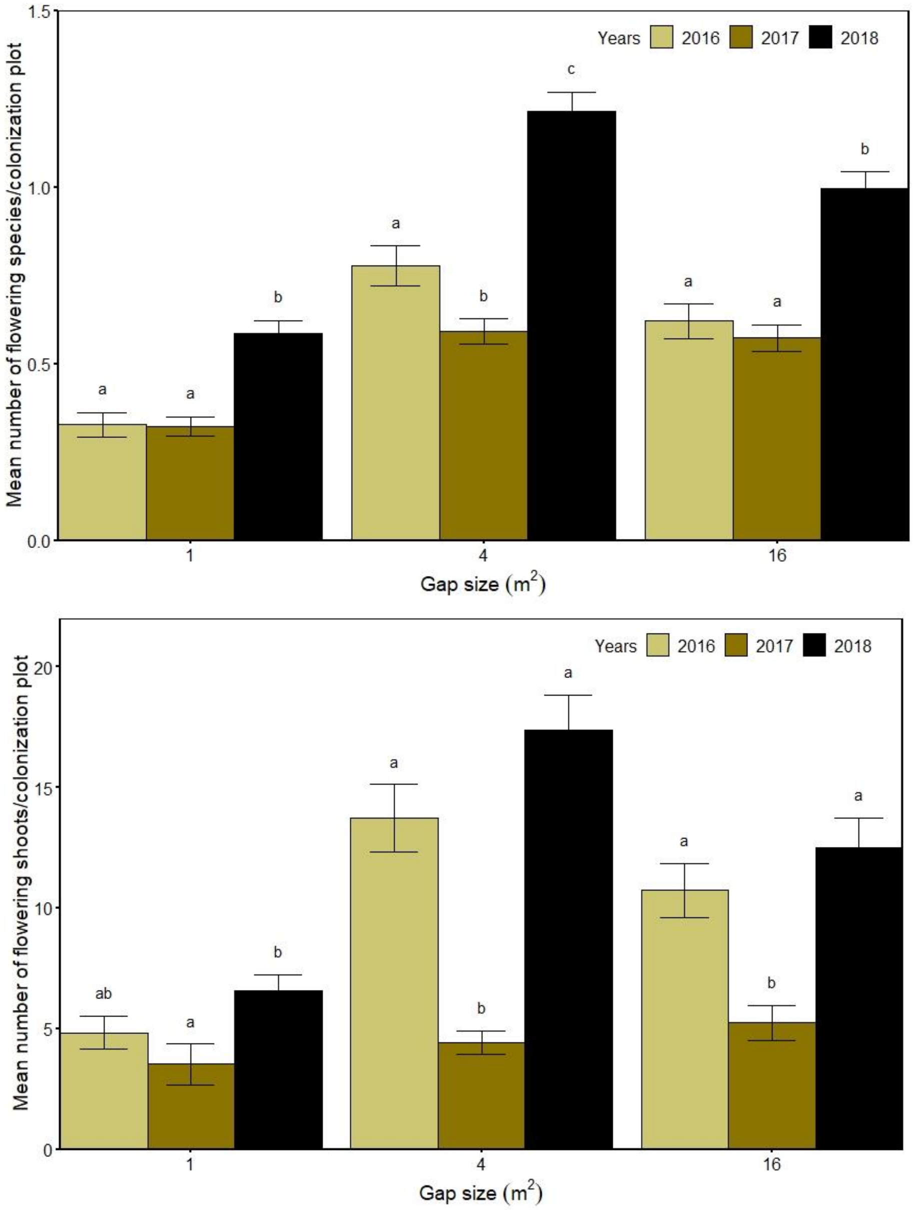


**Figure S3.** Number of flowering sown species (mean ± SE) and number of flowering shoots (mean ± SE) in the 1 $m^2$-sized colonization plots adjacent to different sized establishment gaps (1 $m^2$, 4 $m^2$ and 16 $m^2$) between 2016 and 2018. Letters indicate significant differences between the years within different gap sizes (GLMM and LSD-test, $p < 0.05$).

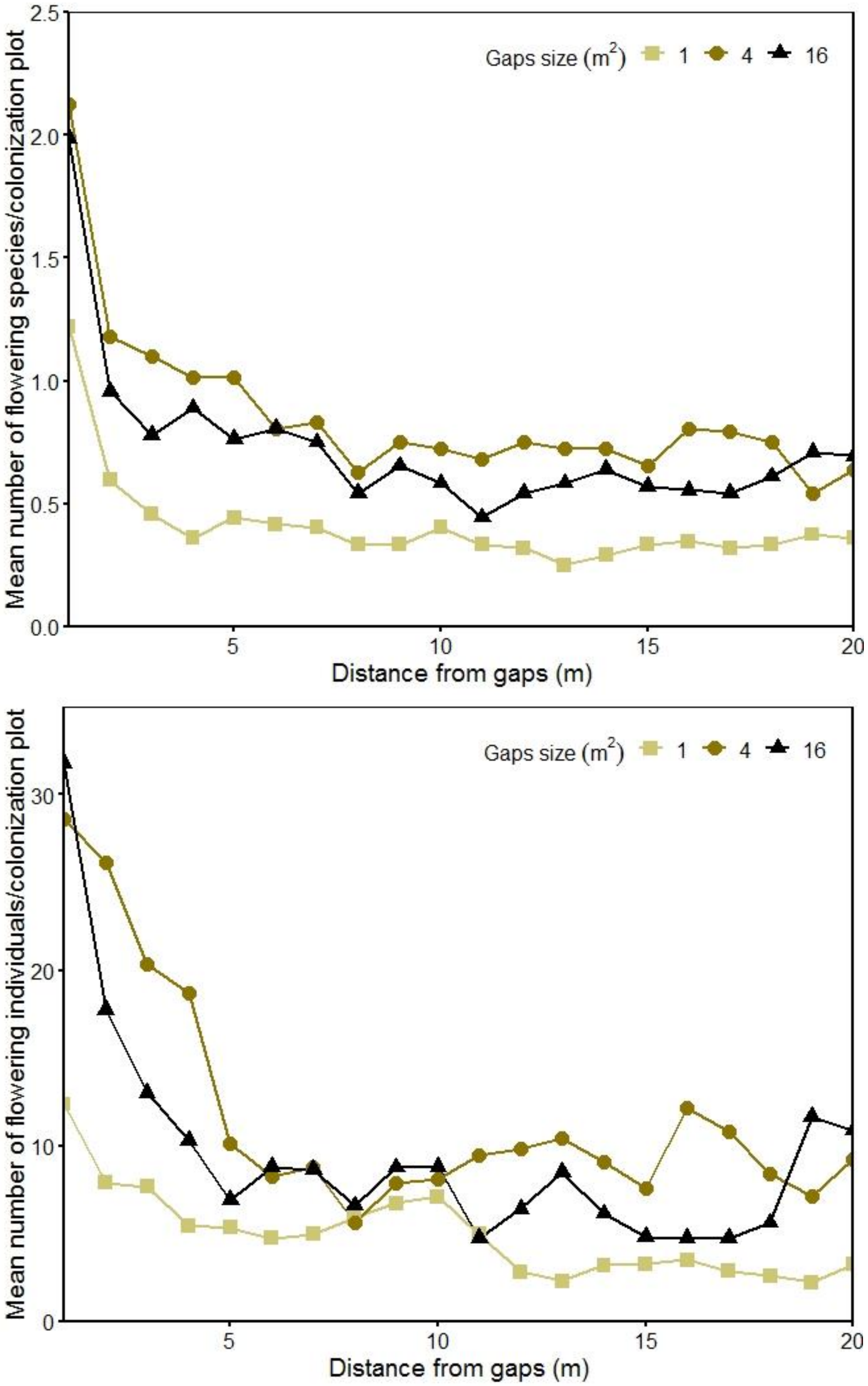


**Figure S4.** Number of flowering species (A) and number of flowering shoots (B) in the 1 m$^2$-sized colonization plots along the 20 m-long transects adjacent to different sized establishment gaps.
